# Disrupting peroxisomes alters lipid metabolism in melanoma and uncovers a novel therapeutic vulnerability in combination with MAPK-targeted therapies

**DOI:** 10.1101/2022.10.25.513718

**Authors:** Fan Huang, Feiyang Cai, Michael S. Dahabieh, Kshemaka Gunawardena, Ali Talebi, Jonas Dehairs, Farah El-Turk, Jae Yeon Park, Christophe Goncalves, Natascha Gagnon, Jie Su, Perrine Gaub, Jean-Sébastien Joyal, John J Mitchell, Johannes V Swinnen, Wilson H. Miller, Sonia V. del Rincón

## Abstract

Melanomas reprogram their metabolism to rapidly adapt to therapy-induced stress conditions, allowing them to persist and ultimately develop resistance. We report that a subpopulation of melanoma cells tolerate MAPK pathway inhibitors (MAPKi) through a concerted metabolic reprogramming mediated by peroxisomes and UDP-glucose ceramide glycosyltransferase (UGCG). Compromising peroxisome biogenesis, by repressing PEX3 expression, potentiates the pro-apoptotic effects of MAPKi via an induction of ceramides, an effect limited by UGCG-mediated ceramide metabolism. Co-targeting PEX3 and UGCG selectively eliminates a subset of metabolically active, drug-tolerant CD36^+^ melanoma persister cells, thereby sensitizing melanoma to MAPKi and delaying resistance. Increased levels of peroxisomal genes and *UGCG* are found in patient-derived MAPKi-relapsed melanomas, and simultaneously inhibiting PEX3 and UGCG restores MAPKi sensitivity in multiple models of therapy resistance. Finally, triple therapy comprised of a newly identified inhibitor of the PEX3-PEX19 interaction, a UGCG inhibitor and a MAPKi demonstrates potent anti-tumor activity in pre-clinical melanoma models, thus representing a promising approach for melanoma treatment.

**Highlights:**

- Inhibiting peroxisome biogenesis uncovers a metabolic vulnerability in melanoma
- CD36^+^ persister melanoma cells tolerate MAPK-targeted therapy through peroxisome/UGCG mediated metabolic rewiring
- Dual blockade of PEX3 and UGCG potentiates melanoma response to MAPK-targeted therapies and restores therapeutic sensitivity in MAPKi-resistant tumors
- NNC 55-0396 is a PEX3-PEX19 binding inhibitor with potent anti-tumor activity in melanoma

## Introduction

Melanoma is the deadliest form of skin cancer that originates from melanocytes. Cutaneous melanomas are classified into four genomic subtypes, including *BRAF*-mutant (∼50%), *RAS*-mutant (∼25%), *NF1*-mutant (∼15%) and triple wild-type melanomas (1). Current treatments for patients with metastatic melanoma include therapies that target the MAPK pathway (i.e., BRAF and MEK inhibitors) and immunotherapies targeting immune checkpoints, such as CTLA-4 and PD-1/PD-L1 axis. However, as with all therapies, MAPK- and immune-targeted treatments have their limitations, often with patients being intrinsically resistant or developing resistance to therapy (2). Expanding the toolbox of treatments, including novel combination therapies, available for patients with melanoma requires a thorough understanding of the underlying biology of this disease.

Melanomas typically exhibit high intratumoral heterogeneity and phenotype plasticity, which ultimately promote MAPKi-resistance, as a combined result of clonal evolution (MAPKi-mediated cell selection) and the emergence of phenotypically and metabolically distinct cell states with survival advantages upon MAPK inhibition (3, 4). Metabolic reprogramming is a mechanism through which subsets of melanoma cells adapt to microenvironmental cues and MAPK-targeted therapy (5-9). PGC1α-mediated mitochondrial oxidative phosphorylation (OXPHOS) and PPARα-mediated fatty acid oxidation (FAO) are two metabolic pathways which have emerged as promising targets to overcome therapy resistance in melanoma (6-9).

Peroxisomes are highly specialized membrane-bound organelles with vital metabolic functions including beta-oxidation of very-long chain fatty acids (VLCFAs), alpha-oxidation of branched-chain fatty acids (BCFA), biosynthesis of C24-bile acids and ether-phospholipids (EPLs), such as plasmalogens, and reactive oxygen species (ROS) metabolism. In mammalian cells, the peroxins PEX3, PEX16, and PEX19 are required for *de novo* biogenesis of peroxisomes (10, 11). Genetic mutations that lead to deregulated peroxisome biogenesis and accumulation of dysfunctional peroxisomes are linked to severe metabolic disorders, such as X-linked adrenoleukodystrophy (X-ALD) and Zellweger syndrome (11, 12). Patients with peroxisomal disorders often exhibit severely impaired lipid metabolism (12). Recent studies suggest that dysregulated sphingolipid metabolism, marked by increased levels of ceramide and altered sphingomyelin abundance, might be a potential biomarker of peroxisomal disorders (13-16), indicating a previously underappreciated role of peroxisomes in the metabolism of sphingolipids. Moreover, emerging evidence highlights the peroxisome as playing significant roles in maintaining human health, with altered peroxisome functions impacting an expanding list of diseases including cancer, diabetes, and neurodegeneration (reviewed in (17)). Although there has been a growing appreciation for the roles of ether-phospholipids (EPLs) and ceramides in melanoma progression, stress response, and drug sensitivity (18, 19), it is surprising that the role of peroxisomes, namely their lipid metabolism, in melanoma response and/or resistance to MAPK inhibition has been underexplored.

Herein, we sought to investigate the role of peroxisome-mediated lipid metabolism in melanoma response and resistance to MAPK inhibition. We chose genetic inhibition of *PEX3* as a mode to disrupt peroxisome biogenesis in a panel of melanoma cells harbouring different genetic driver mutations. A reduction in PEX3 expression in melanoma cells potentiated their response to MAPK inhibition *in vitro* and *in vivo*. Using mouse models and lipidomic analysis, we showed that *PEX3* low-expressing melanoma cell lines relied on the UDP-Glucose Ceramide Glucosyltransferase (UGCG)-mediated ceramide metabolism pathway for survival. Interrogating a previously published single cell RNA-sequencing (scRNA-seq) dataset (20), we identified a CD36^+^ cell state population with dependence on peroxisome/UGCG-mediated metabolic rewiring for MAPKi tolerance. Finally, we identified an inhibitor of PEX3-PEX19 binding and showed that it worked in concert with a UGCG inhibitor to potentiate melanoma response to MAPK-targeted therapy.

## Results

### Compromising peroxisome biogenesis potentiates melanoma cell response to MAPK-targeted therapies

Continuous exposure to MAPK-targeted therapies, such as BRAF and MEK inhibitors, can trigger metabolic reprogramming to promote cell state transitions, known as phenotype plasticity, enabling melanoma cells to survive and persist, ultimately contributing to disease relapse. Given the emerging role of peroxisomes in cancer, we posited that peroxisome biogenesis and lipid metabolism might change in melanoma cells in response to MAPK-targeted therapies. We re-analyzed publicly available RNA-seq data from four studies assessing transcriptomic alterations in patient samples collected before and after MAPK-targeted therapy (21-24). Gene set enrichment analysis (GSEA) revealed that approximately 70% of patients showed an overall induction of peroxisome-associated genes after treatment with BRAF inhibitors (BRAFi) alone or when combined with a MEK inhibitor (MEKi) (Figure 1A, S1A). Conversely, interrogating the same data, less than 50% of patients showed an enriched gene signature involved in oxidative phosphorylation (Figure S1A), previously shown to promote MAPKi tolerance in a subset of melanoma cells (6, 25). These in silico-derived data support a potential role of peroxisomes in MAPKi-driven metabolic rewiring, which allows melanoma cells to escape therapeutic pressure.

**Figure 1.**
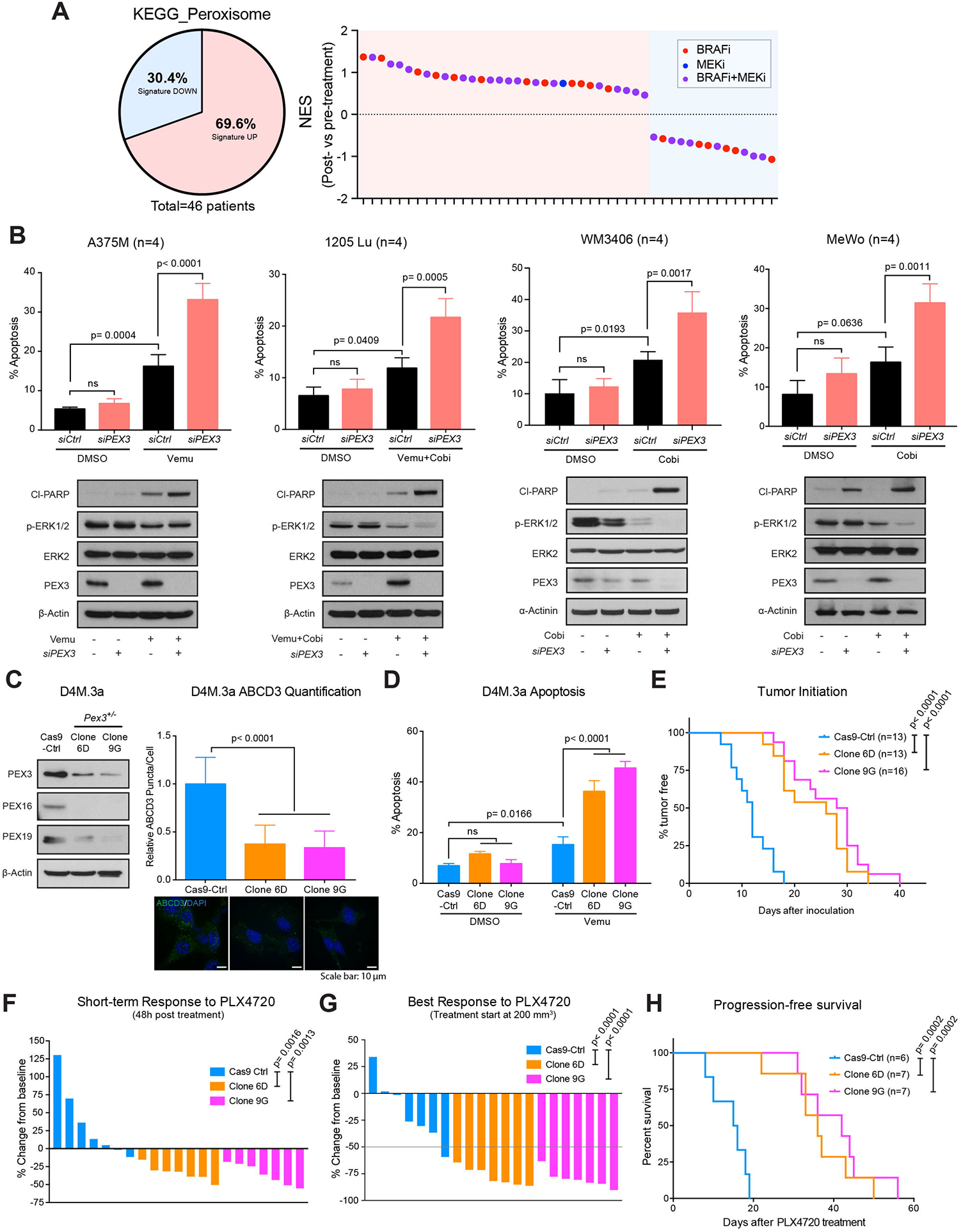
Compromising PEX3 sensitizes melanoma to MAPK inhibition. **(A)** Left: Pie chart showing percentage of patients (n=46) with increased or decreased transcript levels of peroxisome-related genes (KEGG_Peroxisome) after treatment with MAPK-targeted therapies. Right: Normalized enrichment scores (NES) assessing increase (positive) or decrease (negative) of KEGG_Peroxisome gene set in samples from each patient collected post-versus pre-treatment with indicated MAPK inhibitors. **(B)** Top: Percent apoptotic cells as measured by the sum of PI/Annexin V double-positive and Annexin V-positive staining. Bottom: Western blot analysis of the indicated proteins in human melanoma cells following *PEX3* knockdown (or *siCtrl* transfection) and treatment of indicated MAPK-targeted therapy agents (representative of n=4). Equal volumes of DMSO were added to the control groups. Detailed treatment and timeline are presented in Table S1. Two-way ANOVA. **(C)** Western blot analysis of the indicated proteins (left) and relative number of ABCD3 puncta (right) in D4M.3a Cas9-Ctrl, *Pex3*^*+/-*^ Clone 6D and 9G cells. Representative immunofluorescence (IF) staining for ABCD3 (green) and DAPI nuclear stain (blue) are presented (n=3). One-way ANOVA. **(D)** Percent apoptosis (PI^+^/Annexin V^+^, PI^-^/Annexin V^+^) detected in D4M.3a Cas9-Ctrl, 6D and 9G cells following vemurafenib (vemu) or DMSO treatment for 24 hours (n=3). Two-way ANOVA. **(E)** Kaplan-Meier curves showing initiation of D4M.3a Cas9-Ctrl-, 6D- and 9G-derived melanomas. Log-rank test. **(F, G)** Waterfall plots showing **(F)** the short-term response (STR, 48h after treatment initiation) and **(G)** the best response (BR) of D4M.3a Cas9-Ctrl-, 6D- and 9G-derived melanomas to PLX4720. Tumors were allowed to grow to a volume of 200 mm^3^ before PLX4720 treatment started. Values represent % change from baseline. One-way ANOVA. **(H)** Kaplan-Meier curves showing progression-free survival (PFS) of mice bearing D4M.3a Cas9-Ctrl-, 6D- and 9G-derived melanomas, fed with PLX4720 chow. Log-rank test. **(B-D)** Values are represented as mean ± SD. Number of biological replicates is indicated in each graph.

To formally test the impact of disrupting peroxisome biogenesis on response to MAPK-targeted therapy, we knocked down *PEX3* with siRNAs in four human melanoma cell lines harbouring different genetic mutations (BRAF^V600E^-mutant A375M and 1205Lu cells, the NRAS^Q61K^-mutant WM3406 cell line and the NF1^Q1336*^-mutant cell line MeWo) to assess whether this would alter their response to MAPK inhibition. Knockdown of *PEX3* significantly decreased the number of peroxisomes in all four melanoma cell lines (Figure S1B), with no overt effect on cell viability (Figure 1B, S1C). We treated BRAF^V600E^-mutant melanoma cell lines with the BRAF inhibitor vemurafenib (vemu) alone or in combination with the MEK inhibitor cobimetinib (cobi). The non-BRAF-mutant WM3406 and MeWo cells were treated with cobimetinib alone, since the BRAFi would be ineffective in these melanoma subtypes (i.e., *NRAS* and *NF1* mutated). *PEX3* knockdown sensitized all melanoma subtypes to apoptosis induced by the indicated MAPK inhibitors, compared with the same cells transfected with scrambled siRNA control (*siCtrl*) (Figure 1B, S1C). Similar phenotypes were observed following disruption of peroxisome homeostasis by silencing another biogenesis peroxin PEX19 (Figure S1D).

To model our *in vitro* results in mice, we sought to generate an isogenic murine melanoma model with stable *Pex3* knockout. We used CRISPR/Cas9 gene editing to knockout *Pex3* in the BRAF^V600E^-mutant murine melanoma cell line D4M.3a (26). Although none of the 80 clones that we screened showed completed loss of PEX3 expression, we successfully isolated several clones with single allele *Pex3* knockout, two of which we termed *Pex3*^*+/-*^ clone 6D and 9G. As expected, characterization of these *Pex3*^+/-^ cell lines showed decreased levels of PEX3, PEX16, PEX19, and reduced number of peroxisomes (Figure 1C). Similar to our human melanoma cell line data (Figure 1B), the D4M.3a *Pex3*^*+/-*^ clones 6D and 9G were more susceptible to MAPKi-mediated cell death than their Cas9-Ctrl counterparts (Figure 1D). To test the sensitivity of *Pex3*^*+/-*^ melanomas to BRAF inhibition *in vivo*, we next injected D4M.3a Cas9-Ctrl, *Pex3*^*+/-*^ clones 6D or 9G cells into syngeneic mice. A significant delay of tumor onset was observed in mice injected with clone 6D or 9G, compared with mice injected with the Cas9-Ctrl cells (Figure 1E), indicating that compromised peroxisome biogenesis impairs the tumor-initiating ability of melanoma cells. Once initiated, the *Pex3*^*+/-*^ melanomas grew at a similar rate as the Cas9-Ctrl group (Figure S1E). When melanomas of different genotypes reached a volume of approximately 200 mm^3^, mice were administered a diet containing the vemurafenib analog PLX4720 (27). Mice from the 6D and 9G PLX4720 treatment arms showed a rapid decrease in tumor volume within 48 hours of drug administration (short-term response, STR), while the Cas9-Ctrl tumors continued increasing in size during this timeframe (Figure 1F). As the study continued, 6D and 9G treatment groups showed a significantly improved best response (BR) in tumor shrinkage and improved progression-free survival (PFS), compared with the Cas9-Ctrl treatment group (Figure 1G, 1H). The increased treatment benefit in mice whose melanomas were heterozygous for *Pex3* expression was recapitulated in a separate cohort of mice that were administered PLX4720 chow when their tumors reached 800 mm^3^ in size (Figure S1F-S1H).

### Lipidomic analyses of *Pex3*^*+/-*^ D4M.3a melanoma cells reveal altered levels of ceramide-derived lipid species and increased metabolic vulnerability

Next, we investigated potential mechanisms through which inhibiting peroxisomes improves MAPKi response in melanoma. While peroxisome-mediated ROS scavenging is associated with drug tolerance in several tumor types (10, 28, 29), DCFDA staining and N-acetyl-L-cysteine (NAC) scavenger experiments revealed no obvious link between ROS and the apoptotic phenotype observed in *PEX3*-silenced cells upon MAPK inhibition (Figure S2A-S2C). Next, we examined whether mitochondrial integrity, which can be intimately linked to proper peroxisome homeostasis (30), is affected by disrupted peroxisome biogenesis. Analysis of electron micrographs suggested that mitochondrial structure was not affected by PEX3 deficiency in A375M or D4M.3a cells (Figure S2D, S2E). Mitochondrial function was assessed using Seahorse analysis to monitor the oxygen consumption rate (OCR) in our A375M model, and no significant alterations in OCR were observed across any experimental conditions (Figure S2F). These data suggested that compromised peroxisome biogenesis potentiates MAPKi-induced apoptosis in melanoma cells through a mechanism largely independent of alterations in ROS homeostasis and mitochondrial respiration.

Given the essential role of peroxisomes in cellular lipid metabolism (Figure 2A), we next performed lipidomic analyses using our stable *Pex3*^*+/-*^ D4M.3a (6D and 9G) models. We wanted to assess whether alterations in lipid species underpin the anti-tumor phenotypes observed when peroxisome biogenesis is repressed. Our data revealed that *Pex3*^*+/-*^ D4M.3a cells have an increased abundance of phospholipids (PLs), dramatically decreased levels of ether-phospholipids (EPLs), and a slight increase in lysophospholipids (Lyso-PLs) (Figure 2B). We next compared the abundance of each fatty acid-containing species in *Pex3*^*+/-*^ D4M.3a cells versus Cas9-Ctrl cells. Corresponding volcano plots display regions shaded in yellow and blue, which highlight respective increases (≥ 1.5-fold) and decreases (≤ 1.5-fold) in lipid species in *Pex3*^*+/-*^ cells relative to Cas9-Ctrl D4M.3a cells (Figure 2C). Among these altered lipid species, we found 29 commonly upregulated and 69 commonly downregulated species in *Pex3*^*+/-*^ D4M.3a cells (Figure 2D, 2E). Similar to the observed changes in their relative concentrations (Figure 2B), 13 PL species were increased and 60 EPL species were decreased in the *Pex3*^*+/-*^ cells compared with the Cas9-Ctrl cells (Figure 2E). The increased PL and decreased EPL levels in *Pex3*^*+/-*^ D4M.3a cell lines are consistent with clinical phenotypes detected in patients with peroxisomal disorders and corresponding *PEX3* biallelic mutant cell lines *in vitro* (31-33). The latter support the robustness of our lipidomic analyses and the *Pex3*^*+/-*^ D4M.3a cells as a reliable model to understand the biological implications of altering peroxisome biogenesis.

**Figure 2.**
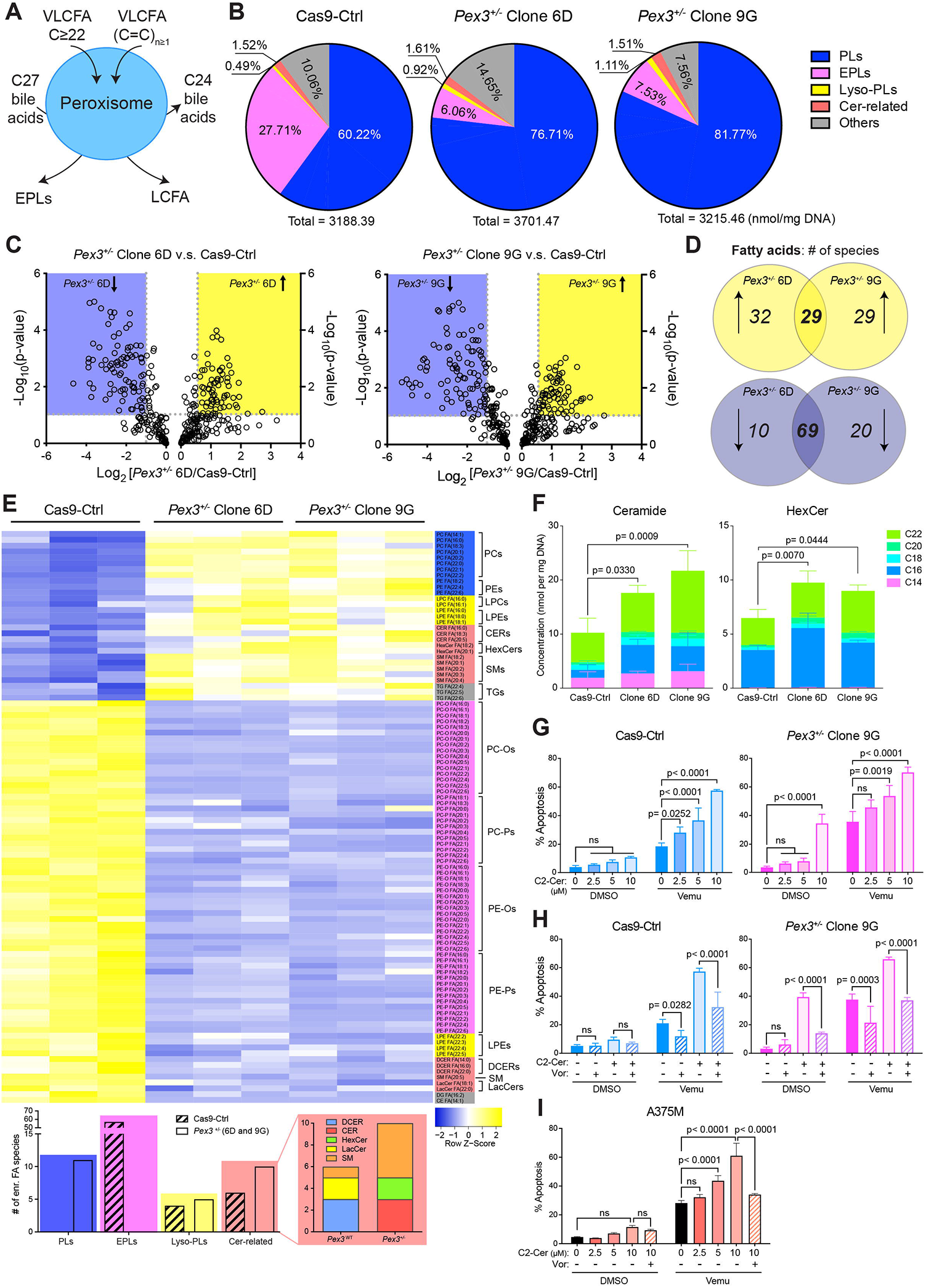
*Pex3*^*+/-*^ D4M.3a melanoma cells have altered lipidomes. **(A)** Schematic of peroxisome-mediated lipid metabolism. **(B)** Pie charts showing lipid composition (relative abundance of each lipid family in % total) in D4M.3a Cas9-Ctrl, Clone 6D and 9G cells. Concentrations of each lipid family (normalized to per mg of DNA) are indicated. (n=3). **(C)** Volcano plots comparing abundance of lipid species in D4M.3a Cas9-Ctrl vs. Clone 6D (left) and D4M.3a Cas9-Ctrl vs. Clone 9G (right). **(D)** Venn diagrams showing lipid species that were significantly increased (top) or decreased (bottom) in D4M.3a Clone 6D and Clone 9G cells, comparing with Cas9-Ctrl cells. **(E)** Top: Heatmap showing lipid species that were commonly altered in D4M.3a *Pex3*^*+/-*^ (6D and 9G) cells comparing with Cas9-Ctrl cells. Bottom: number of lipid species, categorized by family, enriched in D4M.3a Cas9-Ctrl or *Pex3*^*+/-*^ (6D and 9G) cells. **(F)** Concentrations of ceramides (left) and hexosylceramides (HexCer, right) detected in D4M.3a Cas9-Ctrl, 6D and 9G cells (n=3). Two-way ANOVA. **(G-I)** Percent apoptosis (PI^+^/Annexin V^+^, PI^-^/Annexin V^+^) detected in DMSO- or vemu-treated **(G, H)** D4M.3a Cas9-Ctrl (left) or *Pex3*^*+/-*^ Clone 9G (right) or **(I)** A375M cells. Cells were pretreated with **(G, I)** C2-ceramide (C2-Cer) at escalated doses or with **(H, I)** C2-Cer (10μM) and vorinostat (Vor, 1μM) 24 hours prior to vemu treatment (n=3). Two-way ANOVA. All values are represented as mean ± SD.

Interestingly, our lipidomic analysis revealed changes in several sphingolipid species, centering on the synthesis and metabolism of ceramide (Figure 2E-2F, S3). We observed increased levels of ceramides and hexosylceramides (HexCer), which include glucosylceramides (GluCer) and galactosylceramides (GalCer) (Figure 2E, 2F), and decreased levels of dihydroceramide (DCER) and lactosylceramide (LacCer) (Figure S3A). Notably, although several sphingomyelin species (SM) were altered in *Pex3*^*+/-*^ cells (Figure 2E), no consistent changes were observed in total SM level (Figure S3A). Our data are consistent with previous studies showing elevated ceramide levels and altered composition of SM species in patients with peroxisomal disorders (14, 15), further supporting a role of peroxisome as regulators of sphingolipid metabolism (16).

Ceramides are critical mediators of cell fate and lie at the nexus of sphingolipid metabolism (Figure S3B). While increased ceramides can compensate the loss of EPLs due to disrupted peroxisomal function (16), several cell stressors stimulate the production of ceramides to promote apoptosis (34, 35). We hypothesized that the increased susceptibility of melanoma cells with compromised peroxisomes to MAPKi-induced apoptosis is mediated via a mechanism involving ceramide-dependent cell death. We used C2-Ceramide (C2-Cer), a cell-permeable ceramide analog, as a tool to assess its cytotoxicity in Cas9-Ctrl and *Pex3*^*+/-*^ (9G) D4M.3a cells. While Cas9-Ctrl D4M.3a cells were insensitive to C2-Cer induced apoptosis, it potentiated vemu-induced apoptosis in a dose-dependent manner (Figure 2G, left). In *Pex3*^*+/-*^ (9G) cells, which are characterized by increased ceramide levels (Figure 2F), C2-Cer alone was able to induce cell death, albeit at a higher dose (Figure 2G, right), suggesting a potential metabolic vulnerability in peroxisome-compromised cells. When combined with vemu, C2-Cer treatment further induced apoptosis in *Pex3*^*+/-*^ 9G cells (Figure 2G, right). Next, we used the HDAC inhibitor vorinostat (Vor), known to induce peroxisomes (28, 36), to test whether increased peroxisome activity could protect cells from ceramide/vemu-induced apoptosis. As expected, Vor treatment increased peroxisome numbers in both Cas9-Ctrl and *Pex3*^*+/-*^ 9G cells (Figure S3C) and significantly decreased apoptosis induced by C2-Cer and vemu (Figure 2H). Notably, induction of peroxisomes was able protect *Pex3*^*+/-*^ 9G cells against high-dose C2-Cer (Figure 2H, right), further supporting a role of peroxisomes in ceramide metabolism. Similar results were also observed in A375M cells (Figure 2H, S3D). Together, our data suggest that reducing peroxisomes in melanoma cells potentiates their response to MAPK inhibition through their increased ceramide level.

### Dual inhibition of peroxisome biogenesis and UDP-Glucose Ceramide Glucosyltransferase (UGCG) potentiates melanoma response to MAPKi

Having shown that *Pex3*^*+/-*^ cells have a high level of ceramides and apoptose in response to further ceramide increases (Figure 2F, 2G), we next sought to exploit this metabolic vulnerability for therapeutic intervention. *Pex3*^*+/-*^ cells are characterized by increased levels of HexCer (Figure 2F), which include GluCer and GalCer, sphingolipids controlled by the enzymes UGCG, GBA, UGT8, and GALC (Figure S3B). Given that GluCer can be pro-survival in melanoma cells (37), we posited that melanoma cells with compromised peroxisomes rely on the UGCG-catalyzed ceramide-to-GluCer metabolism as a pro-survival mechanism. Dual blockade of PEX3 and UGCG would thus be anticipated to result in a greater clinical benefit. Indeed, knockdown of *Ugcg* significantly increased apoptosis in treatment-naïve *Pex3*^*+/-*^ 9G cells (Figure 3A), indicating that these cells are reliant on UGCG-mediated ceramide clearance for survival. Additionally, knockdown of *Ugcg* further potentiated the response of the *Pex3*^*+/-*^ 9G cells to vemu, resulting in ∼80% cell death within 24 hours of MAPK inhibition (Figure 3A). Consistent with ceramide being pro-apoptotic, *Ugcg*-silenced Cas9-Ctrl D4M.3a cells were also sensitized to vemu-induced apoptosis (Figure 3A). Conversely, silencing *Gba*, which encodes the enzyme catalyzing the reverse GluCer-to-ceramide metabolism, protected *Pex3*^*+/-*^ 9G cells, but not Cas9-Ctrl cells, from vemu-induced apoptosis (Figure 3B). To verify these results in a human melanoma cell system, we generated *PEX3* knockout (*PEX3*-KO) BRAF^V600E^-mutant A375M cells using CRISPR/Cas9 gene editing. Two clones, AG3 and AG7, were isolated and both showed complete knockout of PEX3, decreased levels of PEX16 and PEX19, and dramatically reduced peroxisome numbers (Figure S4A, S4B). Similar to the murine D4M.3a model, genetic silencing of *UGCG* significantly potentiated vemu-induced cell apoptosis in A375M Cas9-Ctrl cells (hereafter referred to as “A375M-Ctrl”) (Figure S4C). Furthermore, knockdown of *UGCG* alone increased apoptosis in both AG3 and AG7 cells, and when these *PEX3*-KO cells were further treated with vemu, a more potent level of apoptosis was observed (Figure S4C).

**Figure 3.**
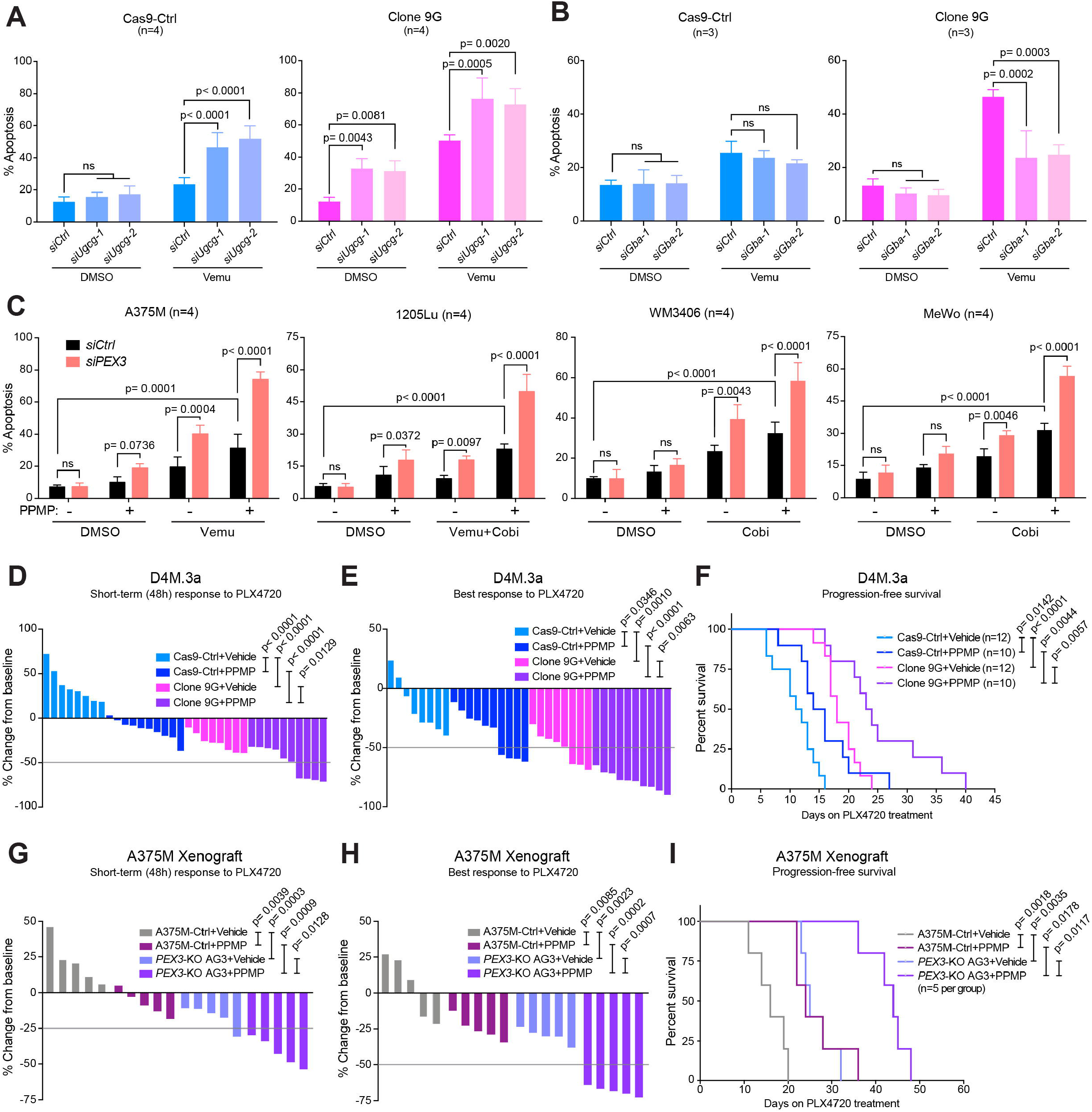
Dual blockade of PEX3 and UGCG sensitized melanoma to MAPK inhibition. **(A, B)** Percent apoptosis (PI^+^/Annexin V^+^, PI^-^/Annexin V^+^) detected in D4M.3a Cas9-Ctrl (left) or *Pex3*^*+/-*^ Clone 9G (right) cells, following **(A)** *Ugcg* or **(B)** *Gba* knockdown and treatment with vemu or DMSO control (n=4 for **A**, n=3 for **B**). **(C)** Percent apoptosis (PI^+^/Annexin V^+^, PI^-^/Annexin V^+^) detected in human melanoma cells following *PEX3* knockdown and treatment with indicated MAPK inhibitors and/or D,L-threo-PPMP (PPMP). Equal volumes of DMSO were added to the control groups (n=4). Detailed treatment and timeline are presented in Table S1. **(A-C)** Values are represented as mean ± SD. **(D, E)** Waterfall plots showing **(D)** the STR (48h after treatment initiation) and **(E)** the BR of D4M.3a Cas9-Ctrl- or 9G-derived melanomas to PLX4720 alone or PLX4720 combined with PPMP. **(F)** Kaplan-Meier curves showing PFS of mice bearing D4M.3a Cas9-Ctrl- or 9G-derived melanomas, treated with PLX4720 alone or PLX4720 combined with PPMP. **(G, H)** Waterfall plots showing **(G)** the short-term (48h) response and **(H)** the BR of A375M-Ctrl- or PEX3*-KO* AG3-derived melanomas to PLX4720 alone or in combination with PPMP. **(I)** Kaplan-Meier curves showing PFS of mice bearing A375M-Ctrl- or PEX3*-KO* AG3-derived melanomas, treated with PLX4720 alone or PLX4720 combined with PPMP. **(D-I)** Number of biological replicates (mice) is indicated in each graph. Detailed experimental timelines are shown in Figure S4F and S4I. **(A-E, G, H)** Two-way ANOVA. **(F, I)** Log-rank test.

To extend our encouraging results with genetic silencing of *UGCG* in our melanoma models, we sought to test the impact of blocking UGCG enzymatic activity on MAPK-targeted therapy response using the inhibitor D,L-threo-PPMP (PPMP), which we show can induce endogenous ceramide level and decrease HexCer (Figure S4D) (38, 39). We first treated *PEX3*-silenced A375M, 1205Lu, WM3406, and MeWo cells with PPMP, and subsequently monitored their response to MAPKi. *PEX3* knockdown combined with PPMP treatment potentiated MAPKi-induced apoptosis in all four melanoma cell lines, compared with PPMP-treated *siCtrl*-transfected cells (Figure 3C). Pharmacological inhibition of UGCG induced apoptosis in both *PEX3*-KO clones AG3 and AG7 clones, but not in A375M-Ctrl cells (Figure S4E). Moreover, PPMP combined with vemu treatment led to the highest level of cell death in the *PEX3*-KO cells (Figure S4E).

We next evaluated the impact of reducing peroxisomes in melanoma cells on their response to combined UGCG inhibition and MAPKi *in vivo*. Briefly, D4M.3a Cas9-Ctrl versus *Pex3*^*+/-*^ (Clone 9G) cells were injected into C57BL/6N mice (Figure S4F), while A375M-Ctrl versus *PEX3*^*-/-*^ (Clone AG3) cells were injected into NOD/SCID mice (Figure S4I). In both models, the peroxisome-deficient (D4M.3a *Pex3*^*+/-*^ 9G and A375M *PEX3*-KO AG3) cells showed delayed tumor initiation compared with their control counterparts (Figure S4G, S4J). When melanomas reached a volume of 200 mm^3^, mice were administered a PLX4720 diet, and PPMP therapy was initiated (Figure S4F, S4I). While both the D4M.3a Cas9-Ctrl- and A375M-Ctrl-derived melanomas continued increasing in size 2 days following PLX4720 monotherapy, the addition of PPMP to PLX4720 rapidly potentiated their short-term response (STR) (Figure 3D, 3G). The long-term treatment with combined PPMP and PLX4720 significantly improved the BR and PFS in the D4M.3a Cas9-Ctrl group, a phenotype that was much stronger in mice bearing *Pex3*^*+/-*^ 9G-derived tumors (Figure 3E, 3F). Similarly, the best BR rate and PFS were observed in the A375M *PEX3*-KO AG3 group when mice were treated with PPMP and PLX4720, compared with all other arms (Figure 3H, 3I). Notably, PPMP treatment, alone or in combination with PLX4720, did not cause any overt toxicity in either mouse model (Figure S4H, S4K). Together, our mouse modelling supports the clinical potential of blocking UGCG and PEX3 to enhance anti-tumor responses to MAPK-targeted therapy.

### Peroxisome and UGCG functions are required for MAPKi-tolerant CD36^+^ persister melanoma cells

Co-targeting PEX3 and UGCG in melanoma not only potentiated the response to MAPK inhibition but also delayed the onset of resistance (i.e., prolonged PFS) in our animal models (Figure 3). We posit that this strategy may eliminate specific populations of MAPKi-persister melanoma cells present in minimal residual disease (MRD), which ultimately lead to the development of drug resistance (20, 40). Using single-cell RNA-seq (scRNA-seq), *Rambow et al*. identified four MAPKi-tolerant melanoma states in a patient-derived xenograft model, termed pigmented, SMC, dedifferentiated (invasive), and neural crest stem cells (NCSC) (20). We re-analyzed the *Rambow* scRNA-seq dataset to examine whether the expression of a peroxisome gene signature (including a panel of peroxisome-associated enzymes and peroxins) was enriched in any of the four MAPKi-tolerant melanoma states. Among the four drug-tolerant states present during the early MAPKi-adapting phase, the peroxisomal gene signature was specifically enriched in the SMC state, which is marked by the expression of the fatty acid transporter cluster of differentiation 36 (CD36) (Figure 4A). Our analysis is consistent with prior work (9) wherein the transcript levels of FAO, which encompassed both mitochondrial and peroxisomal enzymes, were found to characterize the SMC population. Importantly, in our analysis, the expression of the peroxisomal-focussed gene signature and *UGCG* were positively correlated with *CD36* expression in the human melanoma dataset of The Cancer Genome Atlas (TCGA). In contrast, *GBA* and *CD36* expressions were negatively correlated (Figure 4B). Notably, our analysis revealed that *PPARGC1A*, critical for mitochondrial metabolism-mediated MAPKi resistance in a subset of melanoma cells (6, 25), was predominantly expressed in the pigmented MITF^high^ cell state and showed no correlation with *CD36* in the TCGA dataset (Figure 4A, 4B). The CD36^+^ SMC and pigmented melanoma cell states are reliant on altered metabolism for their drug tolerance (8, 25). We thus hypothesized that the CD36^+^ SMC state is distinguished from the mitochondrial-dependent pigmented state by their reliance on a peroxisome and UGCG-dependent mechanism.

**Figure 4.**
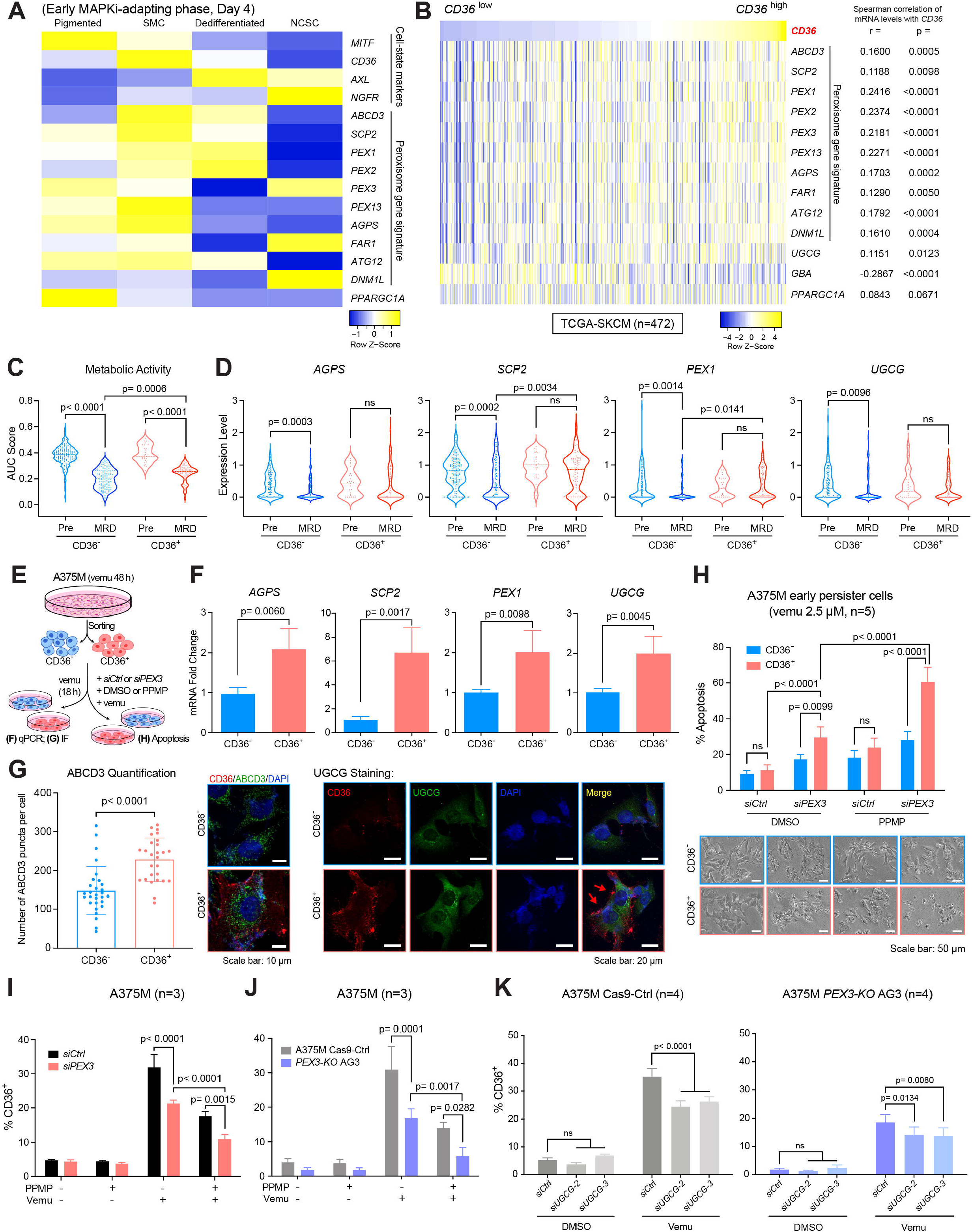
Retained levels of peroxisomes and UGCG are required for survival of CD36^+^ MAPKi-tolerant melanoma cells. **(A)** Heatmap showing relative expression of indicated melanoma cell state-specific markers, a panel of peroxisomal genes, and *PPARGC1A* in 4 drug-tolerant melanoma populations in the MEL006 PDX model during early dabrafenib+trametinib treatment (Day 4). **(B)** Expression of *CD36* and a peroxisomal gene signature, *UGCG, GBA*, and *PPARGC1A* (HTSeq-FPKM) in the GDC TCGA Melanoma dataset (SKCM, n=472). Spearman rank-order. **(C, D)** Violin plots of scRNA-seq data highlighting the distribution of **(C)** a gene signature indicating cancer cell metabolic activity and **(D)** indicated peroxisomal genes and *UGCG* in CD36^-^ (< 2.2) versus CD36^+^ (≥ 2.2) cells, before (Pre) or after dabrafenib+trametinib treatment for 28 days (MRD). **(E)** Schematic of experimental design for **(F-H)**. A375M cells were treatment with vemu (2.5μM) for 48 hours before CD36 staining and subsequent sorting or IF staining. **(F)** Fold change of the indicated mRNAs in vemu-exposed CD36^+^ A375M cells relative to CD36^-^ A375M cells, normalized to *RPLP0* as a reference gene (n=4). The *SCP2* primers amplify the N terminus of the transcript initiated from the proximal promoter, encoding the peroxisome-specific protein SCPx. **(G)** Left: number of ABCD3 puncta in vemu-exposed CD36^+^ versus CD36^-^ A375M cells. Representative IF staining for CD36 (red), ABCD3 (green) and DAPI (blue) are presented. Right: representative IF staining for CD36 (red), UGCG (green) and DAPI (blue) in vemu-exposed CD36^+^ versus CD36^-^ A375M cells (n=3). Red arrows highlight cells stained positive for CD36. **(H)** Percent apoptosis (PI^+^/Annexin V^+^, PI^-^/Annexin V^+^, top) and representative images (bottom) of vemu-exposed CD36^+^ versus CD36^-^ A375M cells following *PEX3* knockdown and/or PPMP treatment (n=5). **(I-K)** Percentage of CD36^+^ populations in **(I)** A375M cells following *PEX3* knockdown and the indicated treatment (n=3), **(J)** A375M-Ctrl versus *PEX3-*KO AG3 cells upon indicated treatment (n=3), or **(K)** A375M-Ctrl versus *PEX3-*KO AG3 cells following *UGCG* knockdown and subsequently treated with vemu (n=4). **(C, D, H-K)** Two-way ANOVA. **(F, G)** Two-sided unpaired t-test. **(F-K)** values are represented as mean ± SD.

We next confirmed that *CD36* marker expression was sufficient to identify the SMC population, which was originally defined by the high expression of a subset of SMC signature genes, including *CD36* (termed “SMC AUCell score”) (20). We examined the *CD36* expression and the SMC AUCell score of these single cells (*Rambow et al*. 2018 dataset) separately. We found that by setting a normalized *CD36* expression ≥ 2.2 as a cutoff (considered as “CD36^+^”) (Figure S5A), we were able to highlight almost identical populations of cells that were defined by SMC AUCell score (≥ 0.05) (Figure S5B). These CD36^+^ cells and SMC populations also showed similar abundance throughout the drug response (Figure S5C). Although dramatic decreases in the cancer cell metabolism gene signature ensue with combined BRAF/MEK inhibition, indicative of an overall low metabolic activity in MRD (20), this metabolic signature was decreased to a significantly lesser extent in the CD36^+^ cells, compared with the CD36^-^ subpopulation (Figure 4C). Consistent with these results, while peroxisome signature genes (*AGPS, SCP2, PEX1*) and *UGCG* expression were significantly decreased in the CD36^-^ cells, the expression of those genes was retained in the CD36^+^ population (Figure 4D). These data suggest a potential mechanistic link between the higher metabolic activity of MAPKi-tolerant CD36^+^ SMC cell state and a role of the peroxisome and UGCG therein. We next formally tested whether peroxisomes and UGCG are required for MAPKi-tolerant CD36^+^ persister cells. Consistent with a previous study (8), treatment with single agent MAPK-targeted therapies vemu, cobi, or combination vemu+cobi increased the percentage of CD36^+^ A375M cells to a similar extent (Figure S5D). We thus used vemu to model the dynamics and drug response/tolerance of these MAPKi-induced CD36^+^ cells. The abundant CD36^+^ population was observed from Day 1 to Day 10 post-vemu treatment (Figure S5E), recapitulating an overall trend observed in the PDX model MEL006 (Figure S5C). After a 48-hour vemu treatment, A375M cells staining negative for both Annexin V and PI were sorted into CD36^-^ and CD36^+^ populations (Figure 4E). As expected, both populations showed lower sensitivity to vemu than the parental A375M cells (Figure S5F). Moreover, CD36^+^ cells were more drug tolerant and developed resistance within a shortest time, compared with either CD36^-^ cells or parental cells (Figure S5F).

Consistent with our scRNA-seq analysis (Figure 4D), vemu-induced CD36^+^ A375M cells showed increased expression of peroxisomal gene signature genes *AGPS, SCP2*, and *PEX1* (Figure 4F), and had an increased number of peroxisomes (Figure 4G, left). Moreover, UGCG expression was higher in the CD36^+^ A375M population, compared with the CD36^-^ A375M cells (Figure 4G, right). Supporting the essential role of the peroxisome in CD36^+^ drug tolerant cells, the CD36^+^ population was more susceptible to apoptosis when *PEX3* was knocked down, compared to the CD36^-^ population. Moreover, this effect was potentiated when UGCG activity was simultaneously blocked using PPMP (Figure 4H). Using both genetic and pharmacological approaches, we further showed that blocking either PEX3 or UGCG was able to decrease the abundance of vemu-induced CD36^+^ A375M cells, and co-targeting both proteins most efficiently eliminated the CD36^+^ population (Figure 4I-4K). Similar results were also observed in the NRAS-driven WM3406 cells upon MEKi treatment (Figure S5G). Together, these data revealed a distinct peroxisome/UGCG-dependent mechanism of tolerance in the MAPKi-induced CD36^+^ melanoma persister cells. Dual blockade of peroxisome biogenesis and UGCG, therefore, enhances efficacy of MAPKi by selectively eliminating these CD36^+^ persister cells.

### Increased peroxisomal activity and UGCG marks MAPKi-resistance and poor outcome in melanoma

Having shown that inhibiting peroxisomes and UGCG can induce death of CD36^+^ MAPKi tolerant cells, we next interrogated the status of CD36, peroxisomes and UGCG in melanomas from patients treated with MAPKi. We re-analyzed a publicly available RNA-seq dataset (21), which included patient melanoma samples collected before, during, and after relapse of MAPKi therapy. In 77% patients (17/22), we observed an overall induction of *CD36* expression following MAPKi treatment (Figure 5A, left). In this patient cohort, expression of the peroxisomal gene signature and *UGCG* expression were most dramatically increased in the relapsed samples (Figure 5A, 5B). Notably, a small group of patients (5/22) showed decreased levels of *CD36* during the course of treatment (Figure S6A, left). No significant pattern of change in peroxisomal gene signature or *UGCG* was observed in this group of samples (Figure S6A, right), suggesting alternative CD36^+^/peroxisome-independent mechanisms through which drug tolerance and resistance can occur. Similar trends were detected in a separate melanoma patient-derived RNA-seq dataset (22, 23). Approximately 76% (19/25) patients showed increased expression of *CD36* following MAPK-targeted therapy (Figure S6B, left). Both peroxisomal genes (*AGPS, SCP2, PEX1*) and *UGCG* were significantly increased after therapy resistance (Figure S6B, right). In this cohort, a slight decrease in *PEX1* and *UGCG* expression was observed in samples collected during therapy, consistent with the scRNA-seq data (Figure 4D).

**Figure 5.**
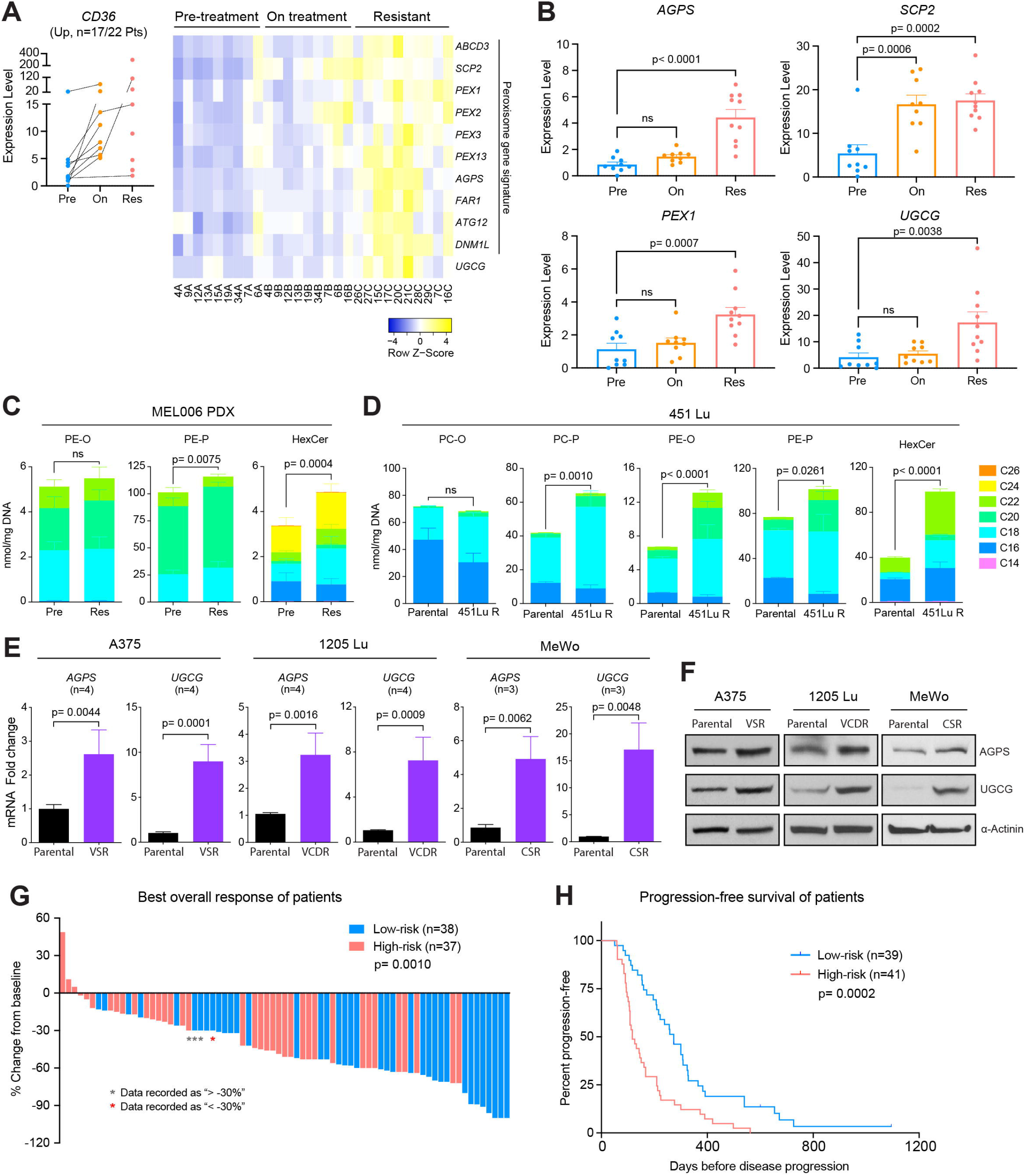
Increased peroxisomal and UGCG activity commonly occurs in melanomas that rapidly acquire resistance to MAPK-targeted therapy. **(A)** Normalized gene expression of *CD36* (left), relative expression of a peroxisomal gene signature (right), and **(B)** expressions of *AGPS, SCP2, PEX1* and *UGCG*, in a cohort of melanoma samples (n=17 out of a total of 22 patients, *Kwong et al*. 2015 dataset) collected pre-, on-, or relapsed on MAPK-targeted therapy showing an overall trend of *CD36* induction upon MAPKi. Values are represented as Mean + SEM. One-way ANOVA. **(C, D)** Normalized abundance of indicated lipid species grouped by carbon chain length detected in **(C)** MEL006 PDX tumor samples collected before or relapsed on dabrafenib+trametinib treatment, or **(D)** 451Lu parental cells versus vemu-resistant 451Lu R cells. Values are represented as mean ± SD. Two-way ANOVA. **(E)** Fold change of *AGPS* and *UGCG* mRNA levels in a panel of MAPKi-resistant melanoma cells relative to their corresponding parental cells, normalized to *ACTB* as a reference gene (n=4). Values are represented as mean ± SD. **(F)** Western blot analysis of the indicated proteins in a panel of parental versus MAPKi-resistant melanoma cells. **(G)** Waterfall plots showing the best overall response and **(H)** Kaplan-Meier curves showing PFS of melanoma patients treated with MAPK-targeted therapy. Risk score ranging between 0-3 was calculated based on expressions of *CD36, AGPS*, and *UGCG* before and after treatment (See Table S5 for detailed information). Patients were subsequently grouped into high-risk (risk score ≥ 2) versus low-risk (risk score ≤ 1). **(E, G)** Two-sided unpaired t-test. **(H)** Log-rank test.

Our data support that peroxisomes and UGCG are essential for the survival of MAPKi-tolerant melanomas and changes in their expression can alter lipid profiles (Figure 2-4). We next interrogated lipidomic data from a human melanoma PDX (MEL006) and cell line model (451Lu) of acquired MAPKi resistance (20, 41) to understand whether these showed lipid profiles indicative of heightened peroxisome and/or UGCG activity. Several ether-phosphatidylethanolamine (PE-P) species were increased in the MEL006 PDX model after tumors relapsed (Figure 5C), strongly indicative of increased peroxisomal/AGPS activity. Similarly, we observed significantly increased abundance of EPL species, including PC-P, PE-O and PE-P, in the vemu-resistant 451Lu R cells, compared with the parental 451Lu cells (Figure 5D). In addition, HexCer was significantly increased in relapsed MEL006 tumors (Figure 5C) and in the 451Lu R cells (Figure 5D). These data are consistent with our results showing that the expression of AGPS and UGCG are upregulated in a panel of MAPKi-resistant melanoma cell lines (Figure 5E, 5F).

Finally, we assessed potential value of *CD36, AGPS* and *UGCG* as biomarkers in predicting patients’ response to MAPK-targeted therapy. We collected data from seven independent studies assessing transcriptomic changes in matched treatment-naïve versus MAPKi-treated melanomas from a total of 80 patients (21-24, 42-44). Using baseline and/or treatment-induced expression of *CD36, AGPS, UGCG* as three individual risk factors, patients were categorized into high-versus low-risk groups (see Methods and Table S5). Indeed, the high-risk group of patients, marked by high or increased expression of *CD36, AGPS* and *UGCG*, showed poorer clinical response to MAPK-targeted therapy and reduced PFS compared to the low-risk group of patients (Figure 5G, 5H).

Together, our data support that increased peroxisomal/AGPS activity and UGCG occur in a significant proportion of relapsed melanomas, which are likely driven through the CD36^+^ drug-tolerant SMC state. Our data further indicate that this subset of melanoma patients may have limited clinical response to MAPK-targeted therapy, which can potentially be improved by combined inhibition of PEX3 and UGCG.

### Overcoming MAPKi resistance by co-targeting peroxisomes and UGCG

We next evaluated the efficacy of dual inhibition of PEX3 and UGCG in a panel of melanoma cell lines with acquired resistance to MAPKi. These cells include previously characterized BRAF^V600E^-mutant A375, WM164 and SK-Mel-28 cells that are resistant to vemu single agent (VSR) (27, 45); newly generated BRAF^V600E^-mutant 1205Lu and SK-Mel-28 cells that are vemu+cobi dual resistant (VCDR); and NF1-mutant MeWo cells that are resistant to cobi single agent (CSR) (See Table S1). Knockdown of either *PEX3* or *UGCG* induced apoptosis in all drug-resistant cells that were cultured in the presence of MAPKi. The UGCG inhibitor PPMP significantly increased apoptosis in *PEX3*-silenced drug-resistant cells, without any such effect in *UGCG*-silenced cells (Figure 6A, S7A), indicating that UGCG inhibition cooperates with *PEX3* inhibition to overcome MAPKi resistance. Similar results were observed in *PEX19*-silenced A375 VSR cells upon PPMP treatment (Figure S7B), suggesting that inhibition of UGCG has potential therapeutic benefit in drug-resistant melanomas with reduced expression of peroxisome biogenesis factors (i.e., PEX3, PEX19).

**Figure 6.**
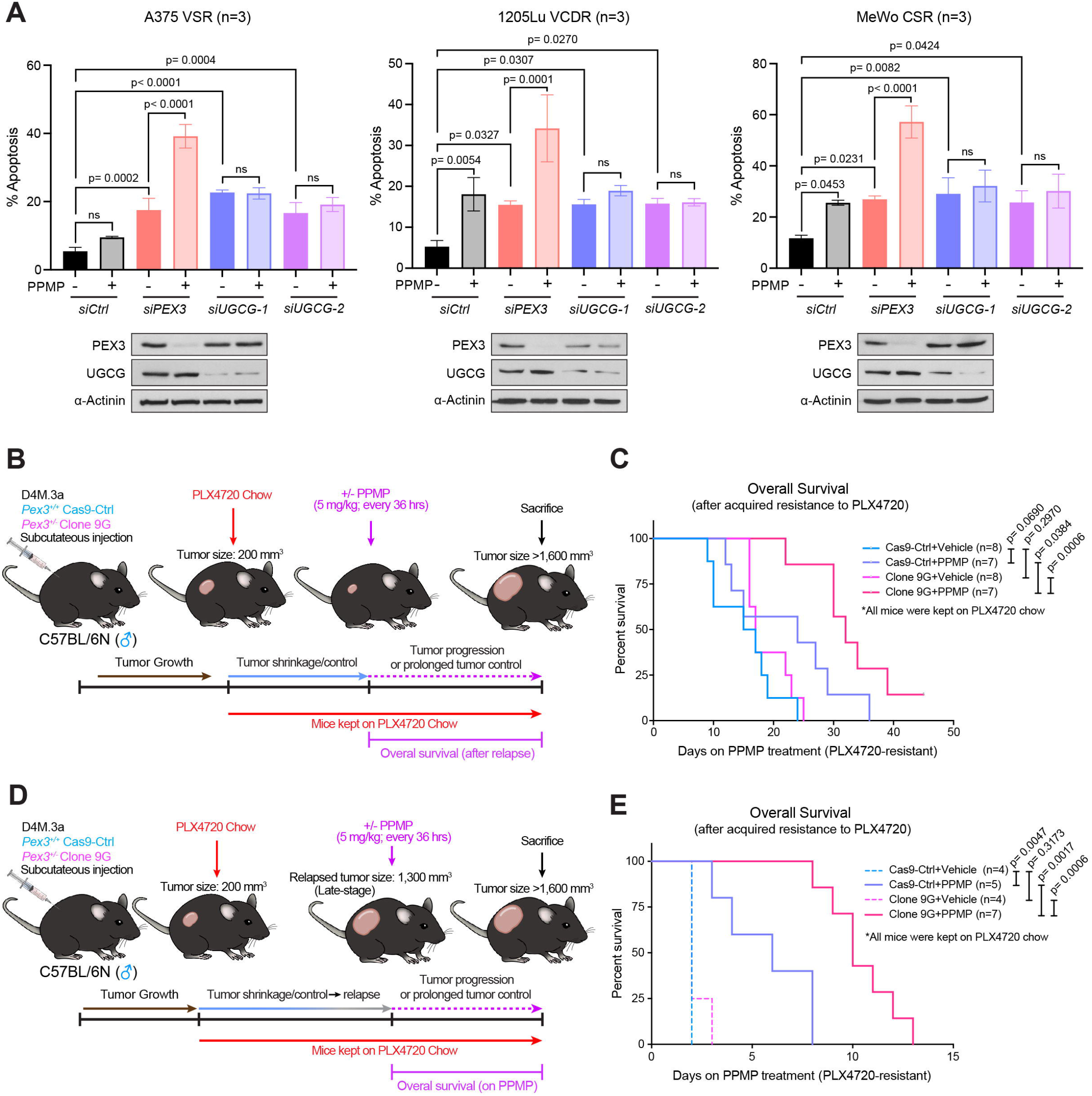
Dual inhibition of PEX3 and UGCG overcomes MAPKi-resistance in melanoma. **(A)** Percent apoptosis (PI^+^/Annexin V^+^, PI^-^/Annexin V^+^) detected in MAPKi-resistant cells following *PEX3* or *UGCG* knockdown and treatment with PPMP. Equal volumes of DMSO were added to the control groups. Cells were maintained in the presence of indicated MAPK inhibitors (Table S1). Detailed treatment and timeline are presented in Table S1. Values represent mean ± SD (n=3). Two-way ANOVA. **(B, D)** Schematic of detailed experimental design. **(C, E)** Kaplan-Meier curves showing overall survival (OS) of Cas9-Ctrl- or 9G-tumor-bearing mice treated with PPMP or vehicle **(C)** after tumors relapsed on PLX4720, or **(E)** after relapsed tumor reached a volume of 1,300 mm^3^. All mice were kept on PLX4720 chow after PLX4720 treatment initiated when individual tumor first reached a volume of 200 mm^3^. Number of biological replicates (mice) is indicated in each graph. Individual tumor growth curves are shown in Figure S7C and S7D, respectively. Log-rank test.

Next, we modeled the *in vivo* efficacy of PPMP therapy in PLX4720-resistant melanoma models that have, or not, altered peroxisomes. Mice bearing size-matched D4M.3a Cas9-Ctrl-versus 9G-derived melanomas (200 mm^3^) were kept on PLX4720 chow until tumor relapse (Figure 6B). PPMP or vehicle treatment was initiated at the time of monitored tumor relapse, annotated with red or grey triangles on the tumor graphs (Figure S7C). Mice were kept on PLX4720 chow and were continually treated with PPMP or vehicle until they were sacrificed due to humane endpoints being reached (Figure 6B). While peroxisome-deficient melanomas (9G cohort) had a delayed onset of PLX4720 resistance compared with Cas9-Ctrl-derived tumors, both groups showed similar growth rates once tumor relapse was detected (Figure S7C, top). No significant difference in overall survival was observed between vehicle-treated Cas9-Ctrl versus 9G groups after acquired resistance to PLX4720 (Figure 6C, S7C). PPMP treatment led to prolonged tumor control in 43% (3/7) of mice in the PLX4720-resistant Cas9-Ctrl group (Figure S7C). However, 100% (7/7) of mice harboring PLX4720-resistant *Pex3*^*+/-*^ melanomas had improved tumor control (Figure S7C) and significantly increased overall survival on the PPMP therapy (Figure 6C). Similar results were observed in another cohort of mice where PPMP was used to treat late-stage therapy-resistant melanomas (Figure 6D, 6E, S7D). Without PPMP treatment, all mice needed to be euthanized within 48 to 72 hours regardless of their *Pex3* status due to their large tumor size (Figure 6E, S7D). Notably, PPMP treatment significantly improved the overall survival of mice bearing Cas9-Ctrl- and 9G-derived late-stage melanomas by approximately 4 and 8 days, respectively (Figure 6E, S7D). We conclude that dual inhibition of PEX3 and UGCG has potential clinical benefit in MAPK-targeted therapy-resistant melanomas.

### The PEX3-PEX19 interaction is druggable with NNC 55-0396 and has therapeutic efficacy in melanoma

Protein-protein interactions are potential vulnerabilities for therapeutic intervention in human disease, including cancer (46). The PEX3-PEX19 interaction is crucial for peroxisome biogenesis (47, 48), and thus provides a potential intervention target with a small molecule inhibitor. A recent screen was conducted to identify compounds which selectively disrupt the PEX3-PEX19 interaction in trypanosomatid parasites without affecting human PEX3-PEX19 binding (49). We leveraged those data to test the top three compounds that the authors identified as disruptors of the human PEX3-PEX19 binding: pregnenolone sulfate (PREGS), 2,3-Dimethoxy-1,4-naphthoquinone (DMNQ), and NNC 55-0396 (NNC) (49). Among these three compounds, NNC, a T-type calcium channel inhibitor (50), significantly decreased peroxisome numbers in human A375M cells (Figure S8A) and was therefore selected for further investigation. Co-immunoprecipitation (co-IP) assay revealed that, NNC dramatically reduced the interaction between PEX3 and PEX19 (Figure 7A). Structural analysis revealed that the predicted binding site of NNC on PEX3 is in close proximity to PEX19-interacting residues (Figure S8B) (51), suggesting a potential mechanism of inhibition through competitive binding.

**Figure 7.**
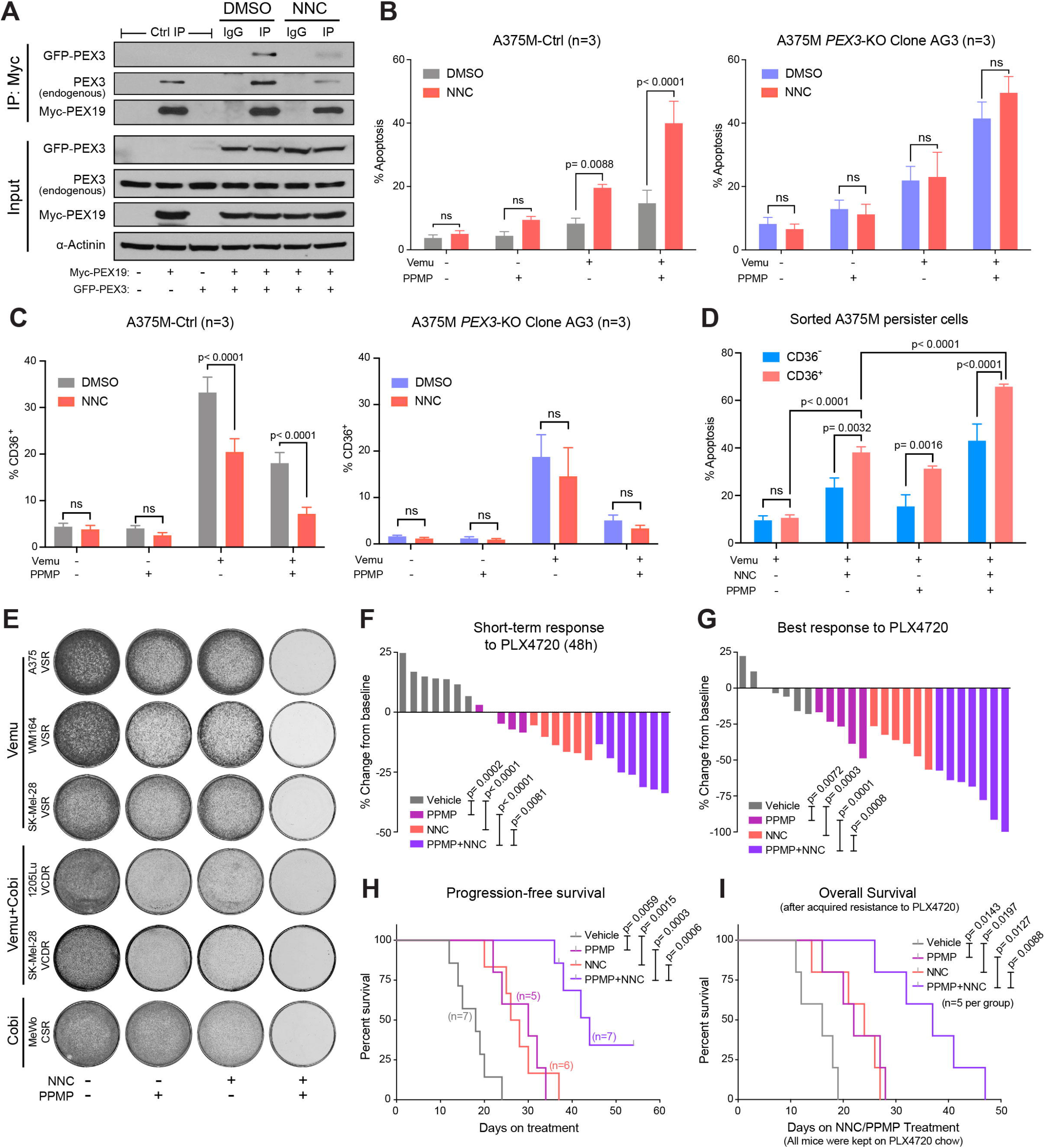
NNC 55-0396 disrupts PEX3-PEX19 interaction and cooperates with PPMP to sensitize melanoma to BRAF inhibition. **(A)** Western blot analysis of PEX3 and PEX19 from Myc co-immunoprecipitation (co-IP) of HEK-293T cells co-transfected with GFP-PEX3- and Myc-PEX19-expressing plasmids, treated with DMSO or NNC 55-0396 (NNC, 4μM, 24h). Left portion displays Myc co-IP of HEK-293T cells transfected with empty vector (EV), Myc-PEX19-, or GFP-PEX3-expressing plasmid alone as negative controls. Immunoblots for inputs (10% of protein from IP) are shown below with α-Actinin probed as loading control (representative of n=3). **(B)** Percent apoptosis (PI^+^/Annexin V^+^, PI^-^/Annexin V^+^) or **(C)** percentage of CD36^+^ populations detected A375M-Ctrl cells (left) or *PEX3*-KO AG3 cells (right) following indicated treatment with NNC (4μM), vemu, and/or PPMP. **(D)** Percent apoptosis (PI^+^/Annexin V^+^, PI^-^/Annexin V^+^) detected in vemu-exposed CD36^+^ versus CD36^-^ A375M cells following NNC and/or PPMP treatment (n=3). **(B-D)** Values represent mean ± SD. **(E)** Colony formation assays of a panel of MAPKi-resistant melanoma cells cultured in the presence of indicated MAPK inhibitors and treated with NNC and/or PPMP for 5 days (representative of n=3). **(F, G)** Waterfall plots showing the **(F)** STR (48h) and **(G)** the BR of A375M-derived melanomas to PLX4720 alone or in combination with NNC, PPMP, or NNC+PPMP. **(H)** Kaplan-Meier curves showing PFS of mice bearing A375M-derived melanomas, fed with PLX4720 chow and simultaneously treated with vehicle, NNC, PPMP, or NNC+PPMP. See detailed experimental design in Figure S8C. **(I)** Kaplan-Meier curves showing OS A375M-tumor-bearing mice treated with vehicle, NNC, PPMP, or NNC+PPMP after relapsed (PLX4720-resistant) tumor reached a volume of 400 mm^3^. All mice were kept on PLX4720 chow after PLX4720 treatment initiated when individual tumor first reached a volume of 200 mm^3^. Detailed experimental design and individual tumor growth curves are shown in Figure S8F and S8G. **(F-I)** Number of biological replicates (mice) is indicated in each graph. **(B-D, F, G)** Two-way ANOVA. **(H, I)** Log-rank test.

Similar to the data obtained in *PEX3*-silenced or -knockout melanoma cells (Figure 1-3), NNC treatment potentiated vemu-induced apoptosis in A375M-Ctrl cells (Figure 7B). The induction of apoptosis was significantly more robust when A375M-Ctrl cells were treated with the triple therapy vemu and NNC in combination with PPMP (Figure 7B). Furthermore, NNC was sufficient to counteract the vemu-mediated increase in CD36-expressing cells (Figure 7C). Combined treatment of NNC and PPMP efficiently and preferably eliminated the vemu-induced CD36^+^ population (Figure 7C, 7D). Similar effects were not observed in NNC-treated A375M *PEX3*-KO (AG3) cells, supporting that the observed biological effects of NNC is through a mechanism involving PEX3 (Figure 7B, 7C). In addition, NNC combined with PPMP efficiently killed a panel of drug-resistant cells cultured in the presence of MAPK inhibitors (Figure 7E), recapitulating a phenotype we observed via genetic inhibition of *PEX3* or *PEX19* (Figure 6A, S7A and S7B).

Finally, we tested *in vivo* efficacy of NNC combined with PPMP and PLX4720 in a preclinical A373M xenograft model. In the first cohort, when tumors reached approximately 200 mm^3^ in size, mice were switched to the PLX4720 diet and simultaneously treated with NNC, PPMP or NNC+PPMP (Figure S8C). NNC administered with PPMP significantly potentiated the PLX4720 anti-tumor response in A375M-derived melanomas, compared with any other groups, a phenotype observed as early as 48 hours following the initiation of therapy (Figure 7F, 7G, S8D). Moreover, long-term administration of the triple therapy (NNC+PPMP+PLX4720) resulted in significantly improved BR and SPF, without causing any overt toxicity (Figure 7G, 7H, S8D and S8E). In a separate cohort to model therapy resistance, mice were switched to the PLX4720 diet when tumors reached approximately 200 mm^3^ in size. NNC, PPMP or NNC+PPMP treatments were subsequently initiated when relapsed tumors reached a size of 400 mm^3^ (Figure S8F). NNC treatment significantly prolonged tumor control and improved overall survival of the mice, an effect that was far more pronounced when NNC was combined with PPMP (Figure 7I, S8G). Together, these data provide a proof-of-concept that combining NNC, a drug able to disrupt the PEX3-PEX19 interaction, with a UGCG inhibitor demonstrates efficacy in sensitizing melanoma to MAPK-targeted therapy.

## Discussion

In this present study, we showed that compromising peroxisome biogenesis by targeting PEX3 (or PEX19) sensitizes melanoma to MAPK pathway inhibition and overcomes melanoma resistance to BRAF and MEK inhibitors, a phenotype that is more robust when UGCG is simultaneously inhibited. Such effects are not limited to melanomas with BRAF^V600^ mutations, but also include MEKi-targetable *RAS*-mutant and *NF1*-mutant melanomas, which broadens its potential clinical application. Mechanistically, we identified two lipid pathways, the peroxisome/AGPS-driven EPL biosynthesis pathway, and the UGCG-catalyzed ceramide-to-GluCer pathway, that collaboratively regulate ceramide homeostasis and mediate drug tolerance in a subset of CD36^+^ persister cells and melanoma cells with acquired MAPKi resistance. Furthermore, we identified NNC 55-0396 as an inhibitor of the PEX3-PEX19 interaction, with its potent anti-tumor effects demonstrated in preclinical melanoma models. This study highlights more broadly, the potential of identifying novel protein-protein interactions in melanoma as potential vulnerabilities for therapeutic intervention.

Melanomas typically exhibit high intratumoral heterogeneity and cancer cell plasticity, ultimately favoring their metastasis and resistance to therapies (3, 4, 20, 40). Metabolic reprogramming represents an important mechanism by which melanoma cells switch between different states and quickly adapt to therapy-induced stress conditions (6, 25). For example, the PAX3-MITF-PGC1α axis promotes MAPKi-tolerance in a subset of melanoma cells through a mitochondrial-mediated metabolic shift (7, 20, 52, 53). Blockade of mitochondrial biogenesis combined with MAPK inhibitors can induce apoptosis in these MITF^high^/pigmented melanoma cells (6). These studies provide evidence that targeting cell state-specific metabolism is a promising strategy to selectively kill drug-tolerant melanoma cell subpopulations. Importantly, our work revealed a distinct metabolic rewiring mechanism harnessed by a different subset of melanoma cells, namely the CD36^+^ SMCs, to mediate their tolerance to MAPK-targeted therapies (Figure 4). The CD36^+^ SMC state is an attractive therapeutic target in melanoma, as it represents a significant subset of drug-tolerant melanoma cells, accounting for approximately 20% to 80% of the population persisting from the early MAPKi response phase to MRD (8, 20). Both CD36^+^ SMC and pigmented melanoma cell states can rely on altered metabolism for their drug tolerance (8, 25). Previous studies suggest that SMCs rely on increased FAO, mediated by PPARα, for their survival during MAPK inhibition (8). Another study identified the peroxisomal enzyme ACOX1, a downstream target of PPARα, as a mediator of drug tolerance generally in melanoma persister cells (9). Although the latter study revealed an enrichment of FAO-related genes in SMCs, including a subset of peroxisomal enzymes, the authors did not explore the role of the PPARα-ACOX1 axis specifically in the CD36^+^ SMC population (9). Notably, the FAO-related genes that were enriched in their persister cells (total of 26 genes) also included 11 mitochondrial-specific enzymes and *PPARGC1A* (encoding PGC1α) (9), a key regulator of mitochondria-dependent metabolic rewiring that functions predominately in a distinct MITF^high^/pigmented melanoma persister cell state (Figure 4A) (7, 20, 52, 53). Using CIBERSORTx, a machine-learning framework that deconvolutes gene expression data from bulk tissues (54), we showed that the enrichment of peroxisomal genes in MAPKi-treated melanoma samples (Figure 1A) is indeed associated with SMC induction (Figure S8H). Approximately 41% (19/46) of patients showed an increase of SMC (>1.5-fold increase) following MAPKi treatment (Figure S8H, left). Interestingly, an enrichment of the KEGG_Peroxisome gene set was observed in ∼95% (18/19) of patients in this group. While within patients with no change or decrease in SMC abundance after MAPK-targeted therapies, only 57% (8/14) and 46% (6/13) showed enriched KEGG_Peroxisome gene signature, respectively (Figure S8H, right). Our data further demonstrated that the CD36^+^ SMCs utilize a peroxisome/UGCG-dependent mechanism for their lipid metabolic rewiring and drug tolerance, distinguishing them from the aforementioned mitochondrial/PGC1α-dependent MITF^high^/pigmented state. Dual blockade of PEX3 and UGCG effectively eliminated CD36^+^ cells (Figure 4H-4K, S5G, 7C and 7D), leading to a dramatic induction of apoptosis in MAPKi-treated melanoma cells and significantly improved response to PLX4720 and PFS in our preclinical models (Figure 3 and 7). These data further support that targeting cell state-specific metabolism is an effective way to potentiate drug response. It is also worth mentioning that, while CD36 is a hallmark of the SMC state, it is not functionally associated with the increased FAO that allows SMCs to survive under the MAPKi-induced metabolic stress (8). For example, knocking out *CD36* did not alter fatty acid uptake or FAO rate upon MAPKi treatment in A375 cells. Using a fatty acid transport protein inhibitor, *Aloia et al*. further showed that CD36 does not function as a fatty acid transporter in MAPKi-induced FAO (8). Therefore, existing inhibitors or neutralizing antibodies against CD36 might not be sufficient to eliminate the SMC population. Together, these data further highlight the significance of combining small molecules that disrupt PEX3-PEX19 binding and UGCG inhibitors to target CD36^+^ persister melanoma cells.

Clinically, combined MAPK-pathway inhibitors can often achieve a fast tumor regression in patients with *BRAF*-mutant melanomas and are hence thought to be particularly beneficial when quick debulking is needed (2). Therefore, in community practice, MAPK-targeted therapy is sometimes offered as first-line treatment to symptomatic patients with late-stage, high-risk, BRAF mutated disease and/or central nervous system (CNS) metastasis to promptly improve their clinical conditions (2). In our preclinical models, combining PLX4720 with PEX3 inhibition and the UGCG inhibitor PPMP achieved an accelerated and more robust tumor regression, compared with PLX4720 monotherapy (Figure 3 and 7). These data suggested that dual blockade of PEX3 and UGCG combined with current MAPK-targeted therapy could potentially benefit patients with late-stage melanomas to achieve a prompt improvement of their clinical condition.

Notably, the use of dual BRAF and MEK inhibition still comes with some clinical challenges. For example, only patients with class 1 *BRAF-*mutant melanomas respond to current targeted therapy agents, and acquired resistance invariably occurs, which may also confer cross-resistance to immunotherapy through MAPKi-driven phenotype switching/dedifferentiation (3, 4). Our data showed that co-targeting PEX3 and UGCG could (a) sensitize *RAS*-mutant and *NF1*-mutant melanoma cells to MEKi-induced apoptosis, (b) delay the onset of acquired resistance *in vivo*, and (c) restore drug sensitivity in resistance/relapsed melanomas (Figure 3, 6 and 7). We further demonstrated that dual blockade of PEX3 and UGCG was able to eliminate MAPKi-induced CD36^+^ persister cells (Figure 4 and S5). Together, these data are consistent with previous observations that not only BRAF inhibitors but also MEK inhibitors induce CD36 expression in melanoma cells (Figure S5D) (8) and that the CD36^+^ SMC state is an important transitory state leading to the emergence of other MAPKi-tolerant states, which ultimately promote acquired resistance (20). Importantly, as undifferentiated and NCSC states are also responsible for MAPKi-induced cross-resistance to immunotherapy (3, 4), targeting these CD36^+^ cells by dual PEX3+UGCG inhibition could also potentially sensitize melanoma to combined immunotherapy and MAPK inhibition and prevent cross-resistance. Such an approach is worth testing in future studies as most patients with advanced *BRAF*-mutant melanoma can be administered both therapies if toxicity is tolerated (2).

We acknowledge that one limitation of our *in vivo* mouse modelling is the use of single agent BRAF inhibitor. However, our data collectively show that our results with single agent BRAFi, would likely extend to dual blockade with BRAFi+MEKi in melanoma. We have shown that in patients, both BRAFi alone or combination therapy of BRAFi+MEKi are able to induce peroxisome-associated genes and SMCs (Figure 1A, S1A, S8H). Moreover, dual blockade of peroxisomes and UGCG potentiate melanoma response to combined BRAFi+MEKi and overcome dual drug resistance *in vitro* (Figure 3C, 6A, 7E and S7A). Another interesting observation we made when generating stable *PEX3* knockout cells for our *in vivo* studies was our inability to generate viable *Pex3* knockouts in the murine D4M.3a cancer cell line, while we had no issue doing so in human A375M cells. Both *Pex3*^*+/-*^ D4M.3a cells and *PEX3*^*-/-*^ A375M cells showed decreased peroxisome numbers and increased sensitivity to combined UGCG and BRAF inhibition (Figure 1, 3, S4). Notably, while EPL deficiency is typically observed in patients with loss-of-function mutations in both alleles of *PEX* genes (10, 11), our data showed that single allele loss of *Pex3* led to significantly decreased levels of EPLs in D4M.3a cells (Figure 2). Given that the anti-tumor phenotypes obtained with both cell models (i.e., heterozygous vs homozygous loss of PEX3) were similar, this suggests that a pharmacologic intervention to reduce peroxisomes, such as NNC 55-0396 used herein, which would presumably never completely ablate peroxisomes, would be anticipated to recapitulate the data we have observed genetically.

It is important to note that melanomas can develop acquired resistance to MAPK-targeted therapies through diverse mechanisms (55, 56). Accordingly, upregulation of CD36 were observed in around 75% (17/22 and 18/25) patient samples following MAPKi treatment (Figure 5A, S6). Approximately 25% (5/22) of the patients showed no CD36 induction, nor clear patterns of peroxisomal gene signature or *UGCG* during their disease progression (Figure S6A), indicating that therapy resistance might have occurred in these patients through mechanisms that are independent of CD36 or peroxisomes. Analysis of publicly available datasets further revealed that increased expression of *CD36* combined with high baseline or increased levels of *AGPS* and *UGCG* marks a group of patients with significantly worse clinical response to MAPK-targeted therapy and shorter PFS (Figure 5G, 5H). Consistent with this observation, sorted CD36^+^ A375M persister cells were more resistant to vemu treatment and developed acquired resistance much faster than CD36^-^ persister cells (Figure S5E). While we did not directly test whether dual blockade of PEX3 and UGCG have efficacy in patients with no CD36 induction, combination of *siPEX3* (or NNC) and PPMP showed limited response in CD36^-^ persister cells (Figure 4H, 7D). Therefore, CD36 upregulation might be a useful early predictive biomarker that can help distinguish responders from non-responders to PEX3+UGCG inhibition.

Finally, peroxisomes have only recently been identified as a potential target for cancer management (10). There are no effective and bioavailable inhibitors of peroxisomes currently available for clinical use. Drug repurposing represents an attractive strategy for identifying new uses (i.e., inhibition of peroxisomes) for existing drugs (57). A recent drug screen aiming to discover inhibitors that disrupt trypanosomatid PEX3-PEX19 binding allowed us to identify NNC 55-0396 as a novel inhibitor of human PEX3-PEX19 interaction (49). NNC was originally developed and is now commonly used as a selective T-type calcium channel blocker, with an IC_50_ of 7μM for inhibition of Ca_v_3.1 T-type channels (50). We showed that NNC effectively disrupted PEX3-PEX19 interaction at 4μM concentration (Figure 7A) and that the anti-tumoral effect of NNC is dependent on peroxisomal functions (Figure 7B, 7C). Moreover, NNC is bioavailable and demonstrate potent anti-melanoma activity in combination with PPMP and PLX4720 in our *in vivo* studies (Figure 7F-7I and S7C-S7G). These data support a novel use of NNC as an PEX3-PEX19 binding inhibitor and provide a proof-of-concept that pharmacological blockade of peroxisomes and UGCG represents a promising strategy for the treatment of melanoma.

Together, our findings opened new avenues for therapeutic intervention in augmenting therapeutic responses to MAPKi and overcoming MAPKi resistance via crippling peroxisome function. This study provided a proof-of-concept that dual blockade of peroxisome biogenesis and UGCG, using pharmacological inhibitors of PEX3-PEX19 binding and UGCG, is an effective strategy to sensitize melanoma to MAPK-targeted therapy.

## Methods

### Mice

Male C57BL/6N mice (6-8 weeks old) were purchased from Charles River Laboratories. Female nonobese diabetic (NOD)/severe combined immunodeficiency (SCID) mice (6-10 weeks old) were kindly gifted by Dr. Moulay Alaoui-Jamali. All mice were randomized before injection. D4M.3a Cas9-Ctrl, *Pex3*^*+/-*^ 6D, and *Pex3*^*+/-*^ 9G cells were prepared in PBS and subcutaneously injected to the right flank of male C57BL/6N mice at 30,000 cells/mouse or 100,000 cells/mouse. A375M Cas9-Ctrl (A375M-Ctrl) and *PEX3*-KO (*PEX3*^*-/-*^) AG3 cells were prepared in PBS and subcutaneously injected to the right flank of female NOD/SCID mice at 1,000,000 cells/mouse. Tumor initiation was determined once palpable tumors were formed. Tumors were then measured in length (L) and width (W). Tumor volumes (V) were calculated based on the formula V=3.1416/6*L*W^2^. For PLX4720 treatment, mice were switched to a special diet containing PLX4720 (AIN-76A rodent diet with 417mg PLX4720/kg, Research Diets, Inc), or the corresponding control chow (AIN-76A, Research Diets, Inc), when tumor volumes reached 200 mm^3^.

For *in vivo* drug treatment, D,L-threo-PPMP (PPMP) (Abcam, ab144023) and NNC 55-0396 (NNC) (TOCRIS Bioscience, 2268) were dissolved in DMSO at concentrations of 20 mg/mL, aliquoted and frozen in - 20 °C. Before each injection, PPMP was freshly diluted in 2 parts of EtOH (70%) and 1 part of 1x PBS to a final concentration of 1.25 mg/mL; NNC was freshly diluted to a final concentration of 4 mg/mL in a solution containing 20% DMSO, 10% PEG400 (Sigma-Aldrich, P3265), and 70% PBS. For the PPMP treatment cohort, mice were kept on PLX4720 chow and were co-treated with PPMP by intraperitoneal injection at 5 mg/kg every 36 hours. Treatments were initiated when 1) the initial tumor volume of PLX4720-naïve mice reaches 200 mm^3^, or 2) tumors had relapsed after PLX4720 treatment (tumor volume increase ≥ 10%), or 3) the relapsed tumors had reached a size of 1300 mm^3^ after PLX4720 treatment. For the NNC treatment cohort, female NOD/SCID mice were subcutaneously injected with 1,000,000 A375M cells/mouse. When the tumor volume reached approximately 200 mm^3^, mice were switched to PLX4720 diet and simultaneously treated with NNC by intraperitoneal injection at 20 mg/kg, and/or with PPMP by intraperitoneal injection at 5 mg/kg every 48 hours.

For the PLX4720-relapsed cohort, mice were kept on PLX4720 diet until tumors relapse. NNC and PPMP treatments were then initiated when relapsed tumors reached a volume of approximately 400 mm^3^. Mice bearing D4M.3a-derived melanomas were sacrificed when the tumor volume reached 1600 mm^3^. Mice bearing A375M-derived melanomas were sacrificed when the tumor volume reached 1000 mm^3^.

### Cells and reagents

The benign melanocyte cell line MelST and the human melanoma cell lines A375M and MeWo were cultured in DMEM media supplemented with 10% FBS and 1x Pen/Strep. The 1205Lu cells and WM3406 cells were cultured in RPMI supplemented with 10% FBS, 1x GlutaMax and 1x Pen/Strep. The WM3406 cell line was a kind gift from Dr. April Rose and the MeWo cell line was a kind gift from Dr. Ian R Watson. The vemu-resistant A375 VSR cells were obtained from Dr. Gideon Bollag. The SK-Mel-28 VSR and WM164 VSR cells were previously generated and described as SK-Mel-28R and WM164R, respectively (45). The vemu/cobi dual resistant 1205Lu VCDR cells were generated by culturing of the parental 1205 Lu cells in elevated doses of vemu and cobi simultaneously. SK-Mel-28 VCDR cells were generated by exposing the vemu-resistant SK-Mel-28 VSR cells sequentially to elevated doses of cobi (with the presence of vemu). The cobi-resistant MeWo CSR cells were generated by culturing of the parental MeWo cells in elevated doses of cobi. All culture conditions and treatment timelines of human melanoma cell lines are listed in Table S1. The murine melanoma D4M.3a cell line was a kind gift from Dr. Brinckerhoff, C. E., and was cultured in Advanced DMEM/F12 media supplemented with 5% FBS, 1x GlutaMax and 1x Pen/Strep.

### Generation of CRISPR cell lines

CRISPR/Cas9-mediated knockout of *PEX3*/*Pex3* in A375M and D4M.3a cells was accomplished using commercially available plasmids. Pre-designed single guide RNAs targeting *PEX3* or *Pex3* were constructed into the pSpCas9 BB-2A-GFP (PX458) vector by the manufacturer (GenScript, Piscataway, NJ, USA). A375M and D4M.3a cells were transfected with either control pSpCas9 BB-2A-GFP plasmid (Cas9-Ctrl) or *PEX3*/*Pex3*-targeting sgRNA/pSpCas9 BB-2A-GFP plasmids. Individual GFP positive clones were sorted into single cells in 96-well plates 48□h after transfection. Single cell clones were expanded and subsequently validated for PEX3 status by western blot and sequencing.

### RNA interference

siRNAs were transfected into cells using Lipofectamine™ RNAiMAX Transfection Reagent (Invitrogen, 13778) following the manufacturer’s instructions. Media were changed the next day after cells were incubated with siRNAs for 18 hours. All cells were harvested between 48h-96h after siRNA transfection. All siRNA sequences are listed in Table S2.

### Immunofluorescence

One coverslip was placed per well in a 24-well plate before cells were seeded in each well. At the time of harvesting, media was aspirated, and cells were washed twice with PBS. Cells were then fixed in 4% formaldehyde diluted in PBS for 20_min at room temperature (RT), then washed three times with PBS. Cell membranes were then permeabilized using 0.2% Triton X-100/PBS for 10□min, then washed three times with PBS. Cells were next blocked with 10% BSA/PBS for 1h at RT, washed once with PBS, and then incubated with indicated primary antibodies diluted in 2% BSA/PBS overnight at 4□°C in a humid chamber. Detailed antibody information is listed in Table S3. Next day, cells were washed three times with PBS at RT, and incubated with secondary antibodies (rabbit 594 and mouse 488 Invitrogen, Carlsbad, CA, USA) for 1h at RT in a humid, dark chamber. Cells were then washed three times with PBS at RT, followed by incubation with 1:1000 DAPI/PBS nuclear stain for 15□min. Cells were washed three times with PBS and then mounted to a glass slide using ProLong gold mounting media (Life Technologies, Carlsbad, CA, USA). Slides were stored in the dark until viewing under a DM IL LED (Leica, Wetzlar, Germany) microscope and images were captured via an Infinity3 (Lumenera, Sarasota, FL, USA) camera. Coloring of images was performed using FIJI software (FIJI, Bethesda, MD, USA).

### Flow cytometry-based assays

Following indicated treatments, cells were trypsinized, centrifuged at 240g for 5□min, and washed twice in PBS. For apoptosis detection of non-fixed cells, Alexa Fluor™ 647-Annexin V (Invitrogen™, A23204) and Propidium Iodide (PI) Staining Solution (BD Biosciences, 556463) were diluted in 1× binding buffer (BD Biosciences, 556454), and subsequently mixed with cells following the manufacturer’s instructions. For DCFDA staining, cells were stained with 5μM H2-DCFDA (ThermoFisher Scientific, D399) for 30□min at 37□°C, followed by Annexin V/PI staining. All flow cytometry experiments were conducted on the LSRFortessa (BD Biosciences).

### Western blotting

Immunoblots were performed as previously described (45). Briefly, cells were lysed with RIPA buffer (150 mmol/L Tris-HCl, pH=7, 150mmol/L NaCl, 1% NP-40, 1% sodium deoxycholate, 0.1% SDS) supplemented with protease and phosphatase inhibitors (Roche). Equal amounts of protein samples were loaded, separated on 10% SDS-PAGE gels, transferred to PVDF membranes, and probed with corresponding antibodies. Detailed antibody information is listed in Table S3.

### Quantitative real-time PCR

Cultured cells were pelleted, and RNA was prepared using the E.Z.N.A. total RNA isolation kit (OMEGA Bio-Tek). RNA concentrations were then quantified using a NanoDrop spectrophotometer (ThermoFisher Scientific) and cDNA was prepared from 1mg of total RNA using iScript cDNA Synthesis Kit (Bio-Rad). Target genes were quantified using the Applied Biosystems 7500 Fast Real-Time PCR System with SYBR Green real-time PCR master mix (Applied Biosystems). Two housekeeping genes were used for each assay. Primers used for qPCR are listed in Table S4.

### Electron microscopy

A375M cells were allowed to grow for 48 hours following siRNA transfection to reach a confluence of 80%. D4M.3a Cas9-Ctrl (*Pex3*^*+/+*^), *Pex3*^*+/-*^ Clones 6D and 9G cells were cultured to 70-80% confluence. Cells were then trypsinized and resuspended in twice the volume of culture media. Cells were initially centrifuged in 15 mL conical centrifuge tubes, and approximately 3 ×10^6^ cells per cell line were washed in 1x PBS, then resuspended, and transferred to 1.5 mL microcentrifuge tubes. Cells were then centrifuged at 100g for 10 minutes, washed twice in 1 mL 1x PBS, followed by aspiration of PBS, retaining cell pellets. Using 2% glutaraldehyde fixation buffer (made by mixing 0.4 ml glutaraldehye 50%, 5 ml 0.2M cacodylate buffer pH7.2 and 4.6 mL distilled water), pellets were fixed for one hour at 4°C. Following fixation, pellets were centrifuged at 100g for 5 minutes, and washed twice with wash/storage buffer (made by mixing equal parts of 0.2M cacodylate buffer pH 7.2 with distilled water). Pellets were then processed and imaged according to previously published methods (58).

### Lipidomics analyses

From three separate passages per cell line, D4M.3a Cas9-Ctrl (P2, P7, P8), *Pex3*^*+/-*^ Clone 6D *(P2, P3, P4)*, and *Pex3*^*+/-*^ Clone 9G *(P3, P4, P5)* cells were cultured to 70-80% confluence, trypsinized, washed twice in PBS, pelleted in a 15 mL conical tube, and flash frozen in surrounding liquid nitrogen. The samples were homogenized and mixed with 0.9□mL MeOH:HCl(1□N) (8:1), 0.8□mL CHCl_3_ and 200□μg/mL of 2,6-di-tert-butyl-4-methylphenol (Sigma, B1378). The organic fractions were evaporated, reconstituted in MeOH/CHCl_3_/NH_4_OH (90:10:1.25) and lipid standards were added (Avanti Polar Lipids). Phospholipids were analyzed by electrospray ionization tandem mass spectrometry (ESI-MS/MS) on a hybrid quadrupole linear ion trap mass spectrometer (4000 QTRAP system, AB SCIEX) equipped with a TriVersa NanoMate robotic nanosource (Advion Biosciences) as described (41). Concentrations of lipid species were normalized to DNA quantity, and relative lipid concentrations were presented as nmol lipid per mg DNA. For the purpose of obtaining statistical information corresponding to fold-changes in lipids between cell lines, data were uploaded onto Perseus (MaxQuant). Values and names of lipid species were numerically and textually characterized. Numerical values were then Log_2_ transformed to obtain a normal numerical distribution pattern within sample types. Zeroes were then imputed using standard settings to fall within the normal curve distribution. Triplicates were grouped according to each cell line, and a multiple-sample one-way ANOVA statistical test was performed to determine whether lipid values differed between cell lines. A post-hoc test was then conducted to determine which cell lines exhibited statistically significant (p ≤ 0.05) differences in Log_2_ transformed lipid abundances. To determine enrichment of species between cell lines, a ≥1.5 fold-change (Log_2_ ≥ 0.585) cut-off and p-value ≤ 0.05 (-Log_10_ ≥ 1.301) was set.

### Cellular and mitochondrial bioenergetics measurements

Bioenergetic profiling was performed according to Agilent Seahorse XF Cell Mito Stress Test Kit Guidebook. Seahorse XF-96 cell culture microplates were first coated with a 50μg/ml working stock of poly-D-lysine and incubated at room temperature for 1-2 hours. 200μl of sterile water per well was added and aspirated from each well, then plates were left to air dry in an incubator without the plate lid. 20,000 A375M cells were transfected with *siCtrl* or *siPEX3* and seeded in each well overnight. The next day, cells were changed to DMSO- or vemu-containing media, then transferred to a 37 °C cell incubator with 20% O_2_ and 5% CO_2_ for approximately 18h. Cells were then washed twice in 1x PBS and incubated with Seahorse XF medium (120μL), and incubated for 1 hour in a non-CO_2_ 37 °C incubator for 1h. XF96 sensor cartridges were placed on top of each well and 20μL, 22μL, and 25μL of oligomycin (final conc.= 2.5μM), Carbonyl cyanide-4 (trifluoromethoxy) phenylhydrazone FCCP (final conc. = 2.0μM), and rotenone/antimycin A (final conc.= 0.5μM), were added to ports A, B, and C respectively. Plates were loaded and analyzed for cellular and mitochondrial bioenergetics according to the guidebook, alongside normalization to cell number.

### PEX3 docking

Docking studies were performed with PEX3 and NNC using Autodock Vina (59) based on the crystal structures of enzymes as deposited in the RCSB Protein Data Bank (https://www.rcsb.org/). NNC was centered at the location of PEX19-binding site on PEX3, and genetic algorithm runs were performed for the ligand and receptor.

### Co-immunoprecipitation

HEK-293T cells were transfected with plasmids overexpressing TurboGFP-tagged PEX3 (OriGene Technologies, RG202031) and Myc-DDK-tagged PEX19 (OriGene Technologies, RC201756) using Lipofectamine 2000 (Invitrogen, 11668-019). The next day, media was changed, and cells were treated with 4μM NNC 55-0396 or DMSO control for 24 hours. Cells were then scraped and lysed with cell lysis buffer (HEPES 25 mM pH 7.5, potassium acetate 115 mM, EDTA 1 mM, 1% NP40) supplemented with protease and phosphatase inhibitors (Roche). Equal amounts of cell lysates were incubated with anti-myc (Cell Signaling Technology, 2276S, IP 1:500) antibody for 1.5h and immunoprecipitated with 25μl of Protein G Dynabeads (ThermoFisher Scientific, 10004D) for 1h at 4□. After washing with cell lysis buffer, the immunocomplexes were analysed by western blotting using anti-PEX3, anti-PEX19, and anti-α-Actinin antibodies as indicated (See Table S3).

### Colony formation assay

MAPKi-resistant melanoma cells were seeded into 6-well plates at 100,000 cells per well in the presence of indicated MAPK inhibitors. The next day, NNC (4μM), PPMP (see Table S1), or DMSO control were added to the indicated wells. Cells were subsequently cultured for 5 days, during which media was changed and drugs were freshly added on day 3. At the end of the assay, cells were fixed with 4% formaldehyde diluted in PBS, stained with 0.5% of crystal violet (Sigma-Aldrich, HT90132) diluted in 70%EtOH and photographed.

### Access and re-analysis of previously published datasets

Normalized single-cell RNA-seq data from the MEL006 PDX model were downloaded from the Gene-Expression Omnibus public database, accession number GSE116237 (20). A total of 674 cells were projected into a two-dimensional space using t-distributed stochastic neighbor embedding (t-SNE). To determine the four established drug-tolerant states and cell metabolic activity, we used the AUCell algorithm based on their characteristic gene signatures as previously described (20). Diapause scores were calculated based on normalized expressions (Z-score) of an embryonic diapause gene signature as previously described (60). *CD36* expression was further examined; and individual cells with a *CD36* expression level ≥ 2.2 were considered as CD36^+^ and were highlighted in the t-SNE map.

Normalized RNA-seq data from patient samples (*Kwong et al*. 2015 dataset) were kindly shared by Dr. Genevieve M. Boland and can be found in the European Genome-phenome Archive (EGA S00001000992) (21). Separate RNA-seq datasets (*Hugo et al., 2015* and *Song et al., 2017* datasets) were downloaded from the Gene-Expression Omnibus public database, accession numbers GSE65186 and GSE75313 (22, 23). A final RNA-seq dataset from a total of 6 patients (*Tirosh et al., 2016*) was directly accessed from the original publication Supplementary Table 9 (24). Normalized and background corrected microarray data (*Kakavand et al., 2017, Long et al., 2014* and *Rizos et al., 2014*) were downloaded from the Gene-Expression Omnibus public database, accession numbers GSE99898, GSE61992 and GSE50509 (42-44).

For Gene Set Enrichment Analysis (GSEA) RNA-seq data from matched pre- and post-treatment samples were analyzed through https://www.gsea-msigdb.org/gsea/index.jsp. Three pre-defined gene sets were assessed: KEGG□Peroxisome, KEGG□Oxidative phosphorylation, and REACTOME□Peroxisomal lipid metabolism. For patient response, PFS, and risk analysis, only patients with gene expression (*CD36, AGPS, UGCG*) data from matched tumor samples that were collected before, on-treatment and/or after relapse were further analyzed and categorized into high- versus low-risk groups using 3 risk factors: RF1, RF2 (a or b), and RF3 (a or b). Briefly, baseline expression of *AGPS* (RF2a) and *UGCG* (RF3a) before treatment and fold change of *CD36* (RF1), *AGPS* (RF2b), *UGCG* (RF3b) expression after treatment (compared to before) were normalized into Z-scores within each dataset. High Z-score of each risk factor values 1 risk score. High- and low-risk groups are defined by a risk score ≥ 2 and ≤ 1, respectively. Detailed risk score and patient information is listed in Table S5. CIBERSORTx was run as described (54) through https://cibersortx.stanford.edu/index.php. Briefly, *Rambow et al., 2018* scRNA-seq data was used to generate the SMC Signature Matrix file (Table S6). Relative SMC fractions in each patient sample were then calculated based on the Signature Matrix file.

Processed quantitative lipidomic data from the MEL006 PDX model and lipidomic data from 451Lu and 451Lu R cells (41) were kindly shared by Dr. Ali Talebi, Dr. Jonas Dehairs and Dr. Johannes V Swinnen.

### Statistics

*In vitro* data were represented as mean ± SD. *In vivo* data were represented as mean ± SEM. Prism software (GraphPad) was used to determine statistical significance of differences. Figure legends specify the statistical analysis used and define error bars. P values are indicated in the figures and p < 0.05 were considered significant.

### Study approval

Animal experiments were conducted according to the regulations established by the Canadian Council of Animal Care, and protocols approved by McGill University Animal Care and Use Committee (#2015-7672).

## Author contributions

FH, SVDR designed research studies. FH, FC, MSD, KG, AT, JD, FET, JYP, and PG conducted experiments. FH, FC, MSD, KG, AT, JD, FET, JYP, CG, NG, JS, and PG acquired data. FH, FC, MSD, KG, AT, JD, FET, JYP, CG, NG, JS, and PG analyzed and interpreted data. FH, MSD, AT, JJM, WHM and SVDR wrote, reviewed, and/or revised the manuscript. JSJ and JVS provided administrative, technical, or material support. WHM and SVDR supervised the study.

## Acknowledgments

This research was funded by the Canadian Institutes of Health Research (CIHR) (grant PJT-162260 to SVDR). FH was endowed by McGill Integrated Cancer Research Training Program graduate studentships. We thank Dr. Genevieve M. Boland (Massachusetts General Hospital, Boston, Massachusetts, USA) for sharing RNA-seq data from patient samples (*Kwong et al*. 2015 dataset). We thank Christian Young, Darleen Element, and Sathyen A. Prabhu for experimental advice and technical supports.

