## Supplementary material for "Disrupting peroxisomes alters lipid metabolism in melanoma and uncovers a novel therapeutic vulnerability in combination with MAPK-targeted therapies": Figure S1-S8 and Table S1-S4

### Supplementary Figures

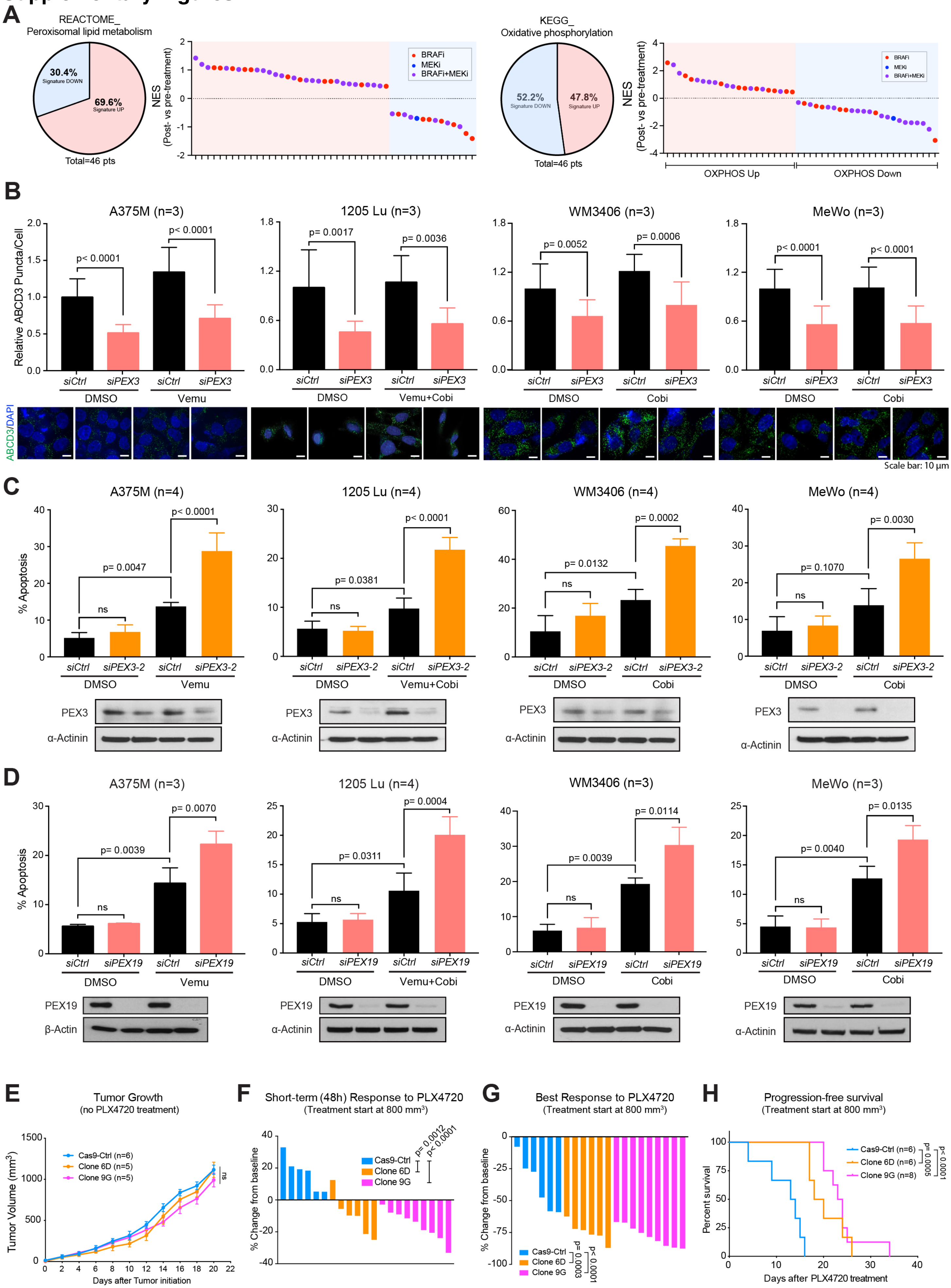

**Figure S1. Compromising peroxisome biogenesis sensitizes melanoma to MAPKi-induced apoptosis.**
**(A)** Pie charts summarizing percentage of patients (n=46) with increased or decreased expressions of indicated gene sets after treatment with MAPK-targeted therapies. Normalized enrichment scores (NES) of samples from each patient (post- versus pre-treatment) are shown. **(B)** Relative number of peroxisomes indicated by ABCD3 puncta in human melanoma cells following *PEX3* knockdown (or *siCtrl* transfection) and treatment of indicated MAPK-targeted therapy agents. Representative immunofluorescence (IF) staining for ABCD3 (green) and DAPI nuclear stain (blue) are presented (n=3). **(C, D)** Percent apoptosis measured as the sum of $PI^+$/Annexin $V^+$ and $PI^-$/Annexin $V^+$ populations (top) and western blot analysis to confirm knockdown (bottom) in human melanoma cells following **(C)** *PEX3* (*siPEX3-2*, see Table S2) or **(D)** *PEX19* knockdown (or *siCtrl* transfection) and treatment of indicated MAPK-targeted therapy agents. **(B-D)** Equal volume of DMSO was added in the control groups. Detailed treatment and timeline are presented in Table S1. Two-way ANOVA. **(E)** Tumor growth curve comparing D4M.3a Cas9-Ctrl-, 6D- and 9G-derived melanomas grown on C57BL6N host mice fed with AIN-76A control diet. Two-way ANOVA. **(F, G)** Waterfall plots showing **(F)** the short-term response (STR, 48h after treatment initiation) and **(G)** the best response (BR) of D4M.3a Cas9-Ctrl-, 6D- and 9G-derived melanomas to PLX4720. One-way ANOVA. **(H)** Kaplan-Meier curves showing progression-free survival (PFS) of mice bearing D4M.3a Cas9-Ctrl-, 6D- and 9G-derived melanomas, fed with PLX4720 chow. Log-rank test. **(F-H)** Tumors were allowed to grow to a volume of 800 $mm^3$ before PLX4720 treatment started. Values are represented as mean ± SD for **(B-D)** and mean ± SEM for **(E)**. Number of biological replicates is indicated in each graph.

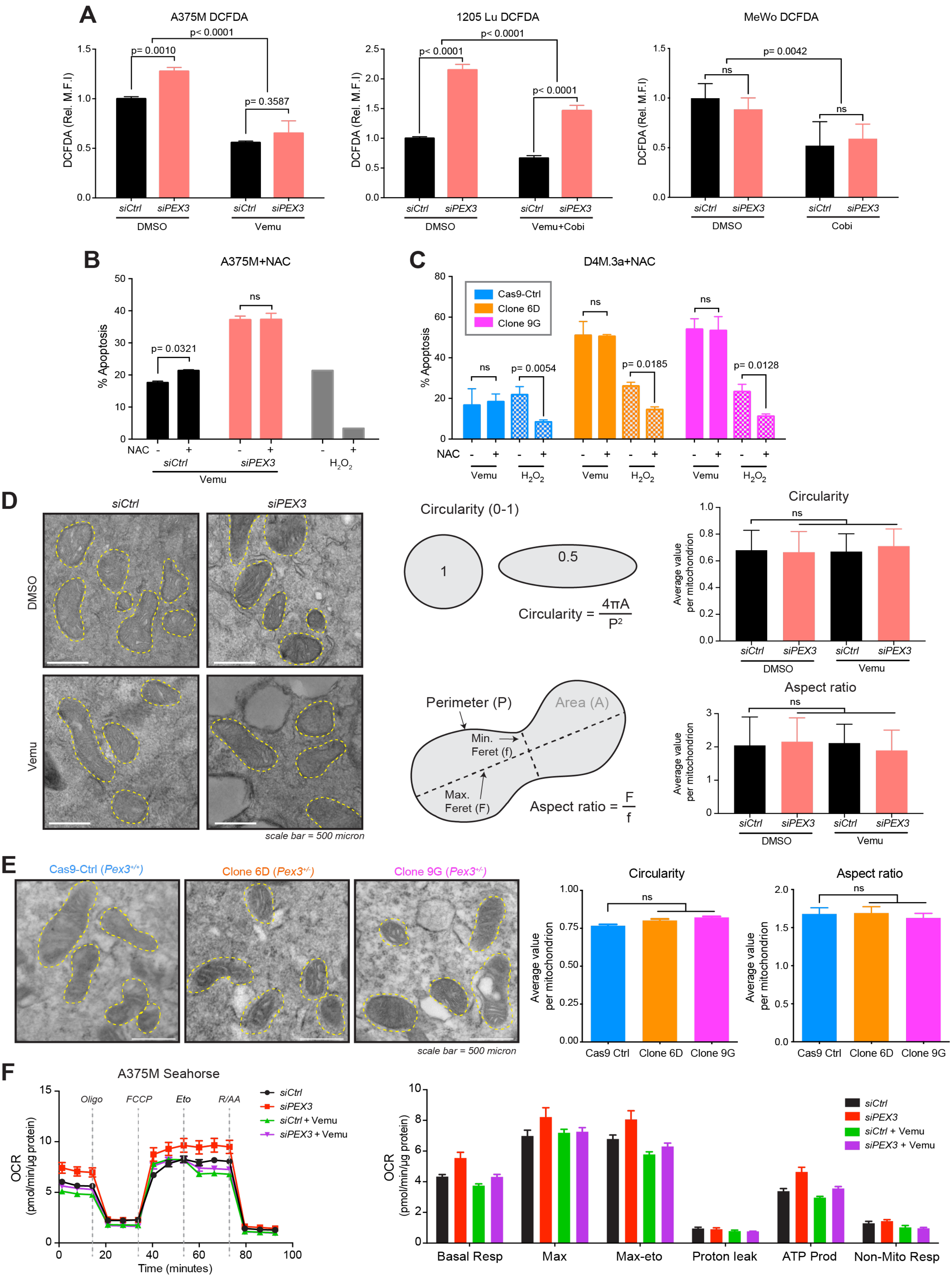
A
A375M DCFDA
1205 Lu DCFDA
MeWo DCFDA
DCFDA (Rel. M.F.I)
p= 0.0010
p< 0.0001
p= 0.3587
p= 0.0042
ns
siCtrl
siPEX3
DMSO
Vemu
Vemu+Cobi
Cobi
B
A375M+NAC
% Apoptosis
p= 0.0321
NAC
Vemu
$H_2O_2$
C
D4M.3a+NAC
Cas9-Ctrl
Clone 6D
Clone 9G
p= 0.0054
p= 0.0185
p= 0.0128
D
Circularity (0-1)
1
0.5
Circularity = $\frac{4\pi A}{P^2}$
Perimeter (P)
Area (A)
Min. Feret (f)
Max. Feret (F)
Aspect ratio = $\frac{F}{f}$
scale bar = 500 micron
Circularity
Aspect ratio
Average value per mitochondrion
E
Cas9-Ctrl ($Pex3^{+/+}$)
Clone 6D ($Pex3^{+/-}$)
Clone 9G ($Pex3^{+/-}$)
Cas9 Ctrl
F
A375M Seahorse
Oligo
FCCP
Eto
R/AA
siCtrl + Vemu
siPEX3 + Vemu
OCR (pmol/min/µg protein)
Time (minutes)
Basal Resp
Max
Max-eto
Proton leak
ATP Prod
Non-Mito Resp

**Figure S2. ROS metabolism and mitochondrial dysfunction do not explain the increased sensitivity of *PEX3*-defficient melanoma cells to MAPK inhibition.**
**(A)** DCFDA relative mean fluorescence intensity (MFI) in human melanoma cells following *PEX3* knockdown and treatment of indicated MAPK-targeted therapy agents (n=3). Equal volume of DMSO was added in the control groups. **(B, C)** Percent apoptosis (PI$^+$/Annexin V$^+$, PI$^-$/Annexin V$^+$) detected in **(B)** *siPEX3*- or *siCtrl*-transfected A375M cells and **(C)** D4M.3a Cas9-Ctrl, 6D and 9G cells, following vemu or DMSO treatment with or without the presence of NAC. Corresponding $H_2O_2$-treated cells were cultured with or without NAC as positive controls (n=3). **(A-C)** Two-way ANOVA. **(D, E)** Representative electron microscope images of **(D**, left**)** *siPEX3*- or *siCtrl*-transfected A375M cells with or without vemu treatment, or **(E**, left**)** D4M.3a Cas9-Ctrl, 6D and 9G cells. The analyses of mitochondrial circularity and aspect ratio in each condition are presented (right). **(D)** Two-way ANOVA. **(E)** One-way ANOVA. **(F)** Seahorse analysis assessing mitochondrial oxygen consumption rate (OCR) in A375M cells following *siPEX3* or *siCtrl* transfection and subsequent vemu or DMSO control treatment (n=3). All values are represented as mean ± SD.

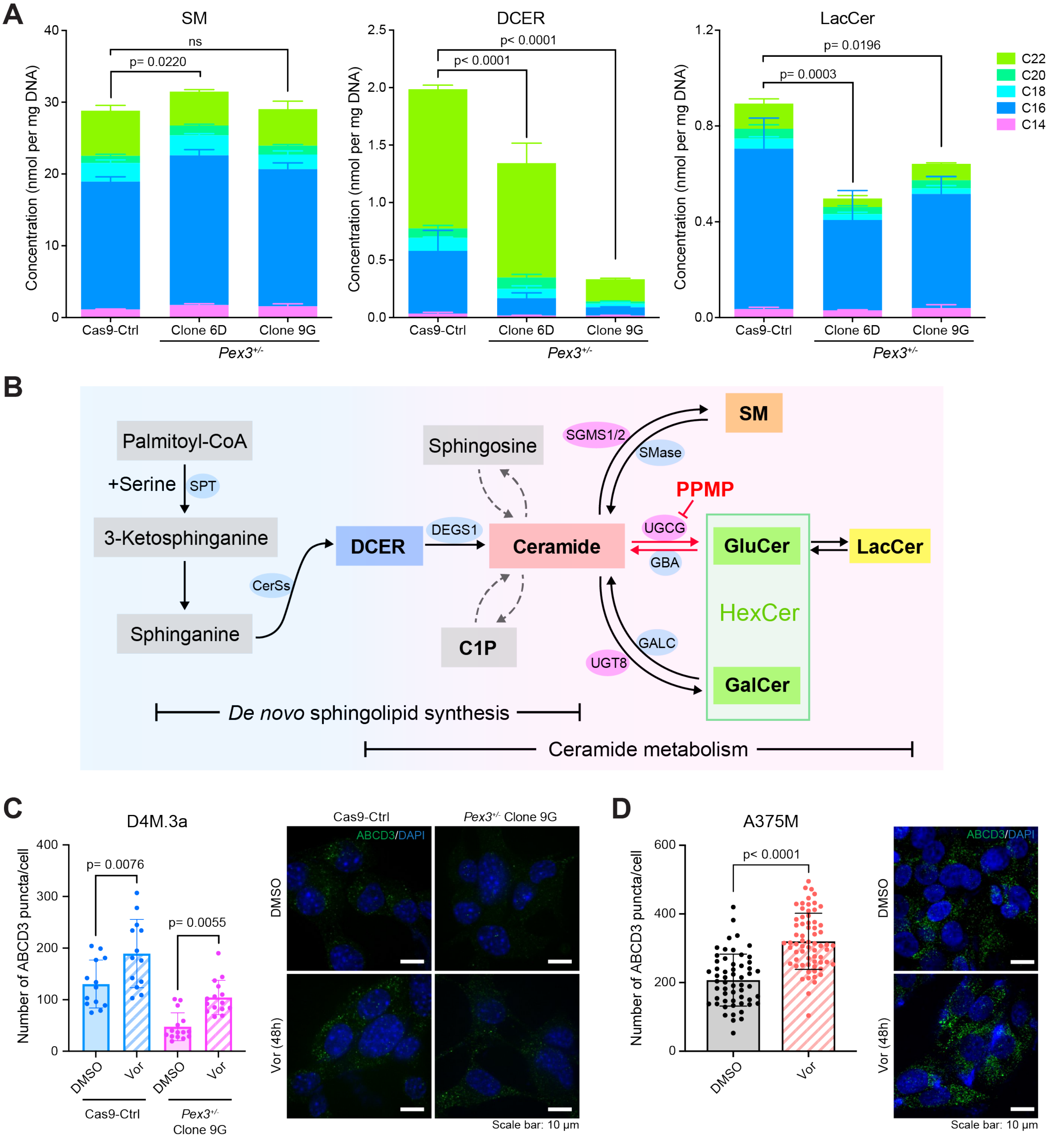


**Figure S3. Single allele knockout of *Pex3* alters lipid composition in D4M.3a melanoma cells.**
**(A)** Concentrations of sphingomyelin (SM), dihydroceramide (DCER), and lactosylceramide (LacCer) detected in D4M.3a Cas9-Ctrl, 6D and 9G cells (n=3). Two-way ANOVA. **(B)** Schematic of the *de novo* sphingolipid synthesis pathway (left) and pathways centering on ceramide metabolism (right). D,L-threo-PPMP (PPMP) blocks UGCG mediated ceramide-to-GluCer metabolism and thereby increases ceramide abundance. **(C, D)** Number of ABCD3 puncta in **(C)** D4M.3a (Cas9-Ctrl versus *Pex3*$^{+/-}$ Clone 9G) cells, or **(D)** A375M cells, treated with DMSO control or vorinostat (Vor, 1μM) for 48 hours. Representative IF staining for ABCD3 (green) and DAPI nuclear stain (blue) are presented. **(C)** Two-way ANOVA. **(D)** Two-sided unpaired t-test. All values are represented as mean ± SD.

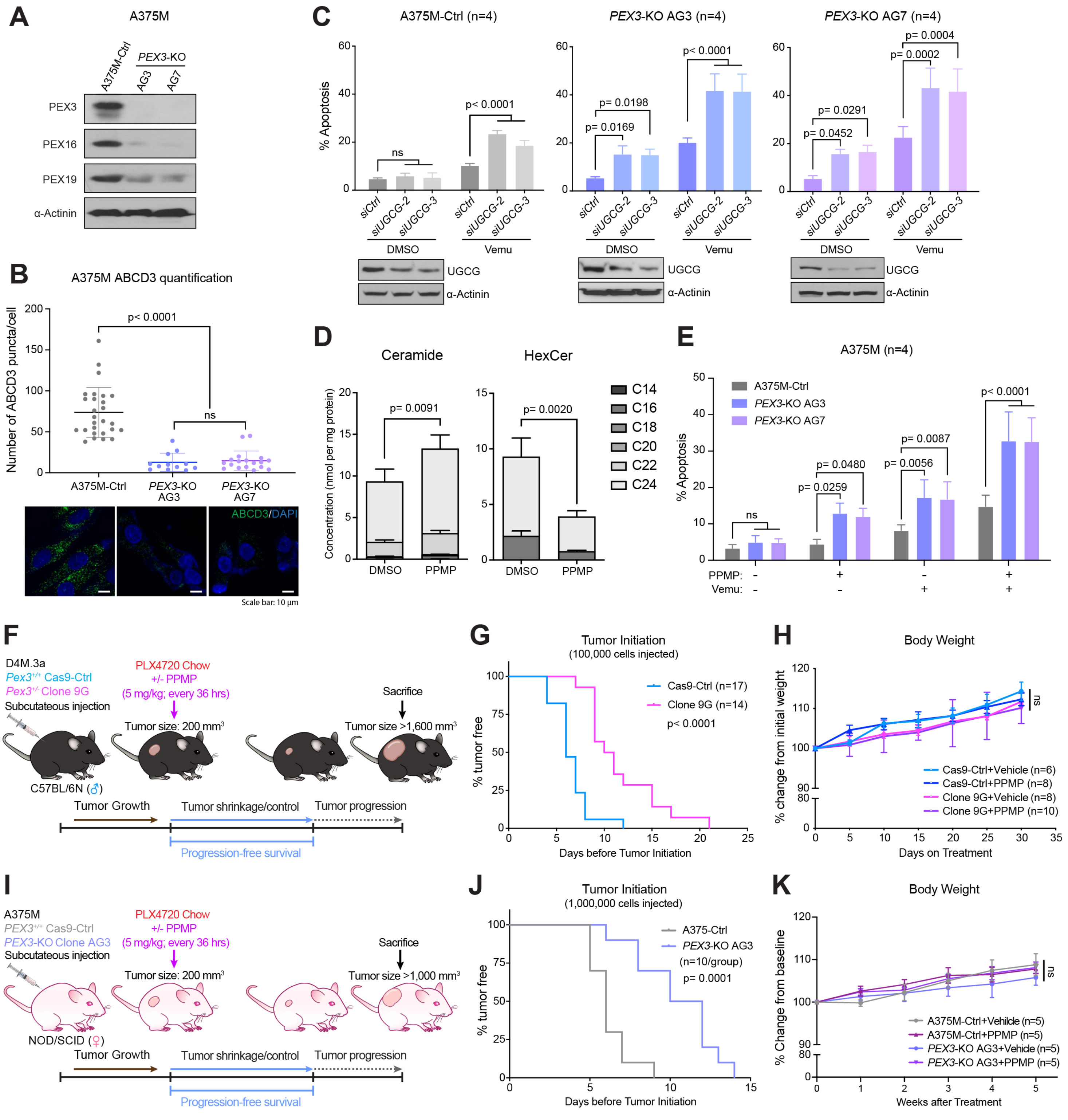


**Figure S4. UGCG blockade increases vemu sensitivity in PEX3-deficient melanomas.**
**(A)** Western blot analysis of the indicated proteins and **(B)** number of ABCD3 puncta in A375M Cas9-Ctrl (A375M-Ctrl), *PEX3*-KO (*PEX3*$^{-/-}$) Clone AG3 and AG7 cells. Representative IF staining for ABCD3 (green) and DAPI nuclear stain (blue) are presented (n=3). One-way ANOVA. **(C, E)** Percent apoptosis (PI$^{+}$/Annexin V$^{+}$, PI$^{-}$/Annexin V$^{+}$) detected in A375M-Ctrl, *PEX3*-KO AG3 and AG7 cells following **(C)** *UGCG* knockdown and treatment with vemu or DMSO control (n=4), or **(E)** PPMP treatment alone or in combination with vemu (n=3). **(D)** Concentrations of ceramide and HexCer detected in A375M cells treated with PPMP (7.5$\mu$M) or DMSO for 24 hours (n=3). **(C-E)** Two-way ANOVA. **(B-E)** Values are represented as mean ± SD. **(E, H)** Schematic of the experimental design, related to Figure 3. **(F, I)** Kaplan-Meier curves showing initiation of **(F)** D4M.3a Cas9-Ctrl- or 9G-derived melanomas (100,000 cells per mice injected), and **(I)** A375M-Ctrl or AG3 (*PEX3*-KO)-derived melanomas (1,000,000 cells per mice injected). Log-rank test. **(G, J)** Relative weight change (% initial body weight prior to treatment) of mice bearing **(G)** D4M.3a Cas9-Ctrl- or 9G-derived melanomas, or **(J)** A375M-Ctrl or AG3 (*PEX3*-KO)-derived melanomas after indicated treatment. Number of biological replicates (mice) is indicated in each graph. Two-way ANOVA. **(G, J)** Data represented as mean ± SEM.

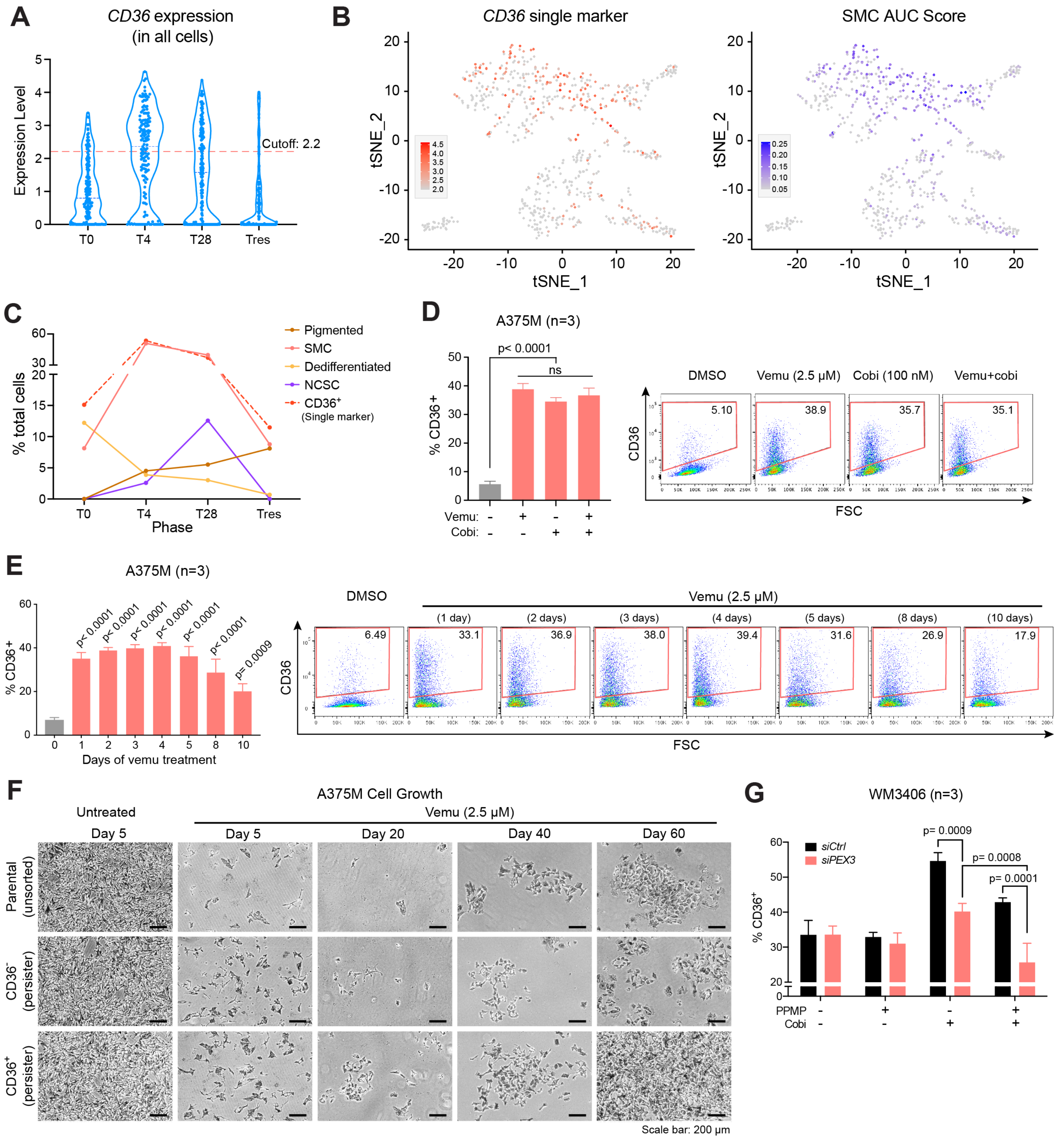


**Figure S5. CD36 marks a distinct population of melanoma persister cells.**
**(A)** Violin plot of scRNA-seq data highlighting the distribution of *CD36* and the cutoff line set to distinguish CD36$^-$ (< 2.2) and CD36$^+$ (≥ 2.2) cells in different phases of MAPKi treatments. **(B)** A total of 674 melanoma cells (*Rambow et al.* 2018 dataset) were projected in a two-dimensional space by t-SNE, comparing the distribution of CD36$^+$ cells (left) defined by high *CD36* expression as a single marker (normalized gene expression ≥ 2.2, colored in red) versus SMCs (right) defined by AUCell analysis (SMC AUCell score ≥ 0.05, colored in purple). **(C)** Dynamics of the different melanoma cell states at the indicated time points. **(C, D)** Percentage of CD36$^+$ populations in A375M cells following **(C)** indicated MAPK inhibitors treatment for 48 hours, or **(D)** vemu treatment for indicated time. Representative flow cytometry imagines are shown for each timepoint (n=3).

One-way ANOVA. **(F)** Representative image of parental (unsorted), CD36⁻ and CD36⁺ persister A375M cells (sorted after 48 hours treatment with vemu) following vemu treatment for indicated days (representative of n=4). **(G)** Percentage of CD36⁺ populations in WM3406 cells following *PEX3* knockdown and the indicated treatment (n=3). Two-way ANOVA. Data represented as mean ± SD.

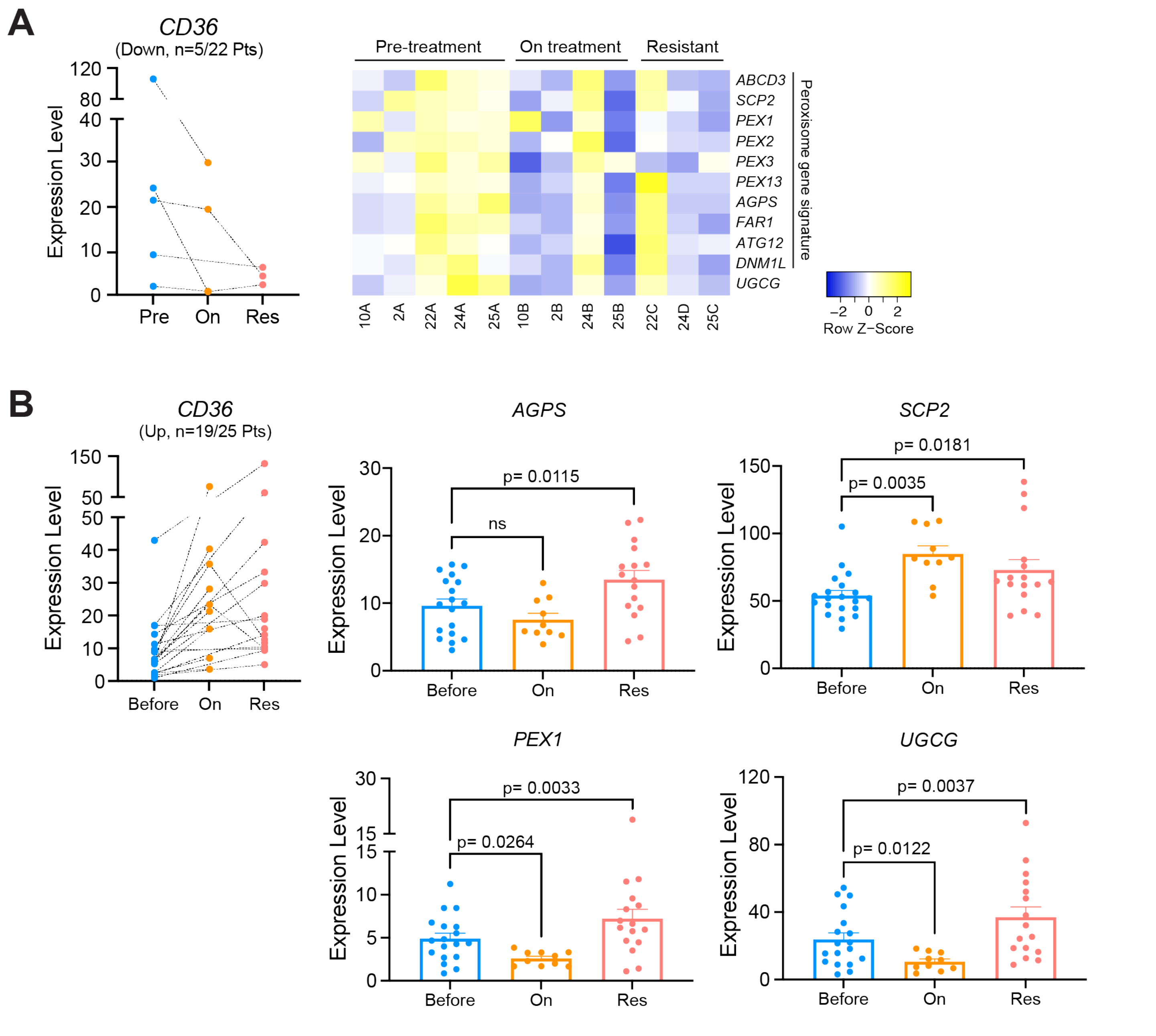


**Figure S6. MAPKi-induced upregulation of *CD36* is associated with increased peroxisome and UGCG activity in therapy-resistant melanomas.**

**(A)** Expression of *CD36* (left) and relative expression of a peroxisomal gene signature and *UGCG* (right) in a cohort of melanoma samples collected pre-, on-, or relapsed on MAPK-targeted therapy. n=5 out of a total of 22 patients (*Kwong et al.* 2015 dataset), with an overall trend of *CD36* downregulation following MAPKi treatment. **(B)** Expression of *CD36* (left) and expressions of *AGPS*, *SCP2*, *PEX1* and *UGCG* (right), in a cohort of melanoma samples (n=19 out of a total of 25 patients, *Hugo et al., 2015* and *Song et al., 2017* datasets) collected pre-, on-, or relapsed on MAPK-targeted therapy showing an overall trend of *CD36* induction upon MAPKi. Values are represented as Mean + SEM. One-way ANOVA.

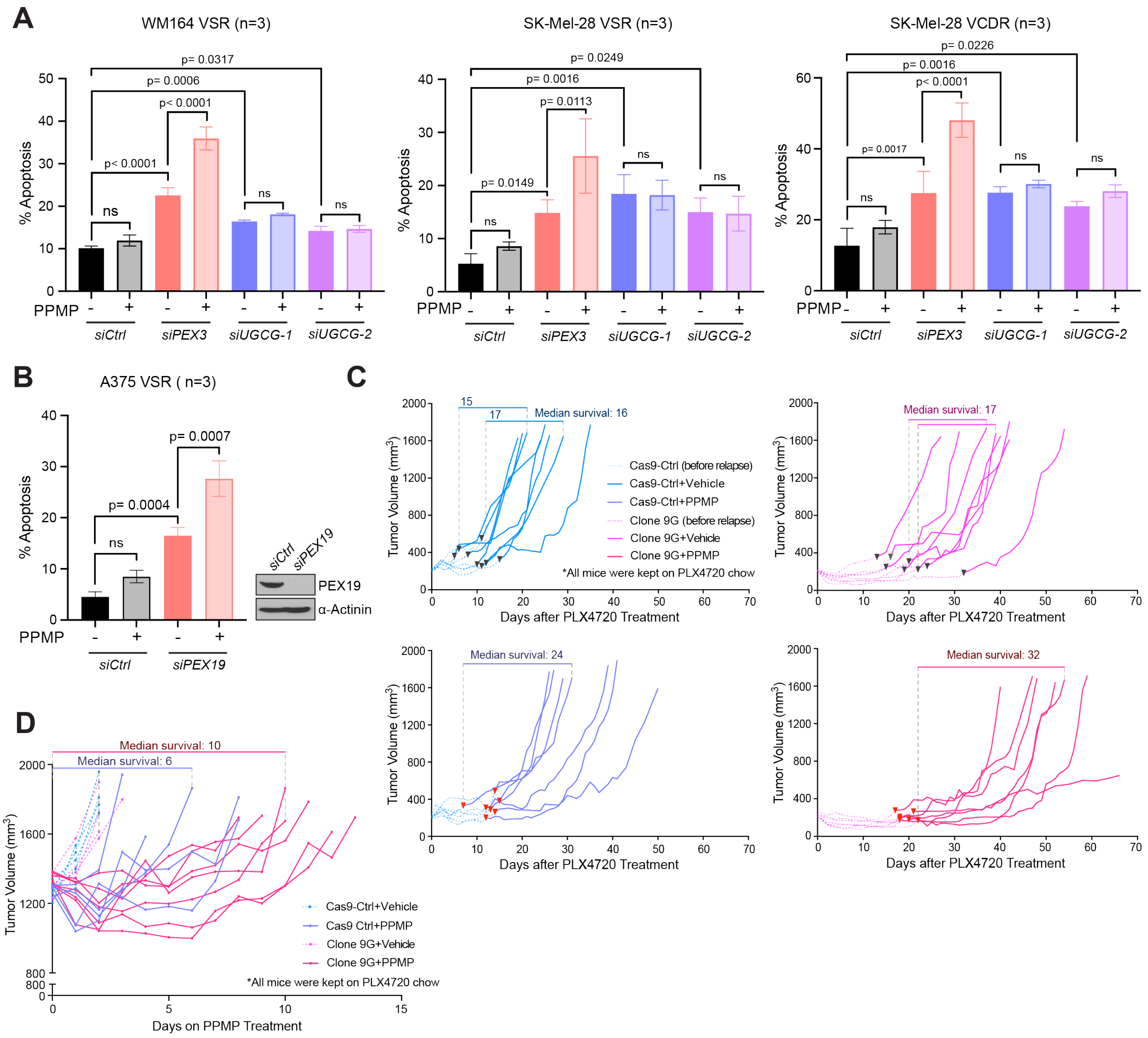


**Figure S7. MAPKi-resistant melanomas are sensitive to combined inhibition of peroxisomes and UGCG.**
**(A, B)** Percent apoptosis ($PI^+$/Annexin $V^+$, $PI^-$/Annexin $V^+$) detected in **(A)** a panel of MAPKi-resistant melanoma cells following *PEX3* or *UGCG* knockdown, or **(B)** A375 VSR cells following *PEX19* knockdown. Cells were maintained in the presence of indicated MAPK inhibitors (Table S1) and were treated with PPMP or equal volume of DMSO as control (Detailed treatment and timeline are presented in Table S1). Values represent mean ± SD (n=3). Two-way ANOVA. **(C, D)** Individual growth of D4M.3a Cas9-Ctrl- or 9G-derived melanomas in mice receiving indicated treatment, related to Figure 6C and 6E, respectively. Mice were treated with PPMP or vehicle **(C)** after tumors relapsed on PLX4720, or **(D)** after relapsed tumor reached a volume of 1,300 $mm^3$. Median survival is indicated in each graph. All mice were kept on PLX4720 chow after PLX4720 treatment initiated when individual tumor first reached a volume of 200 $mm^3$. Number of biological replicates (mice) is indicated in each graph.

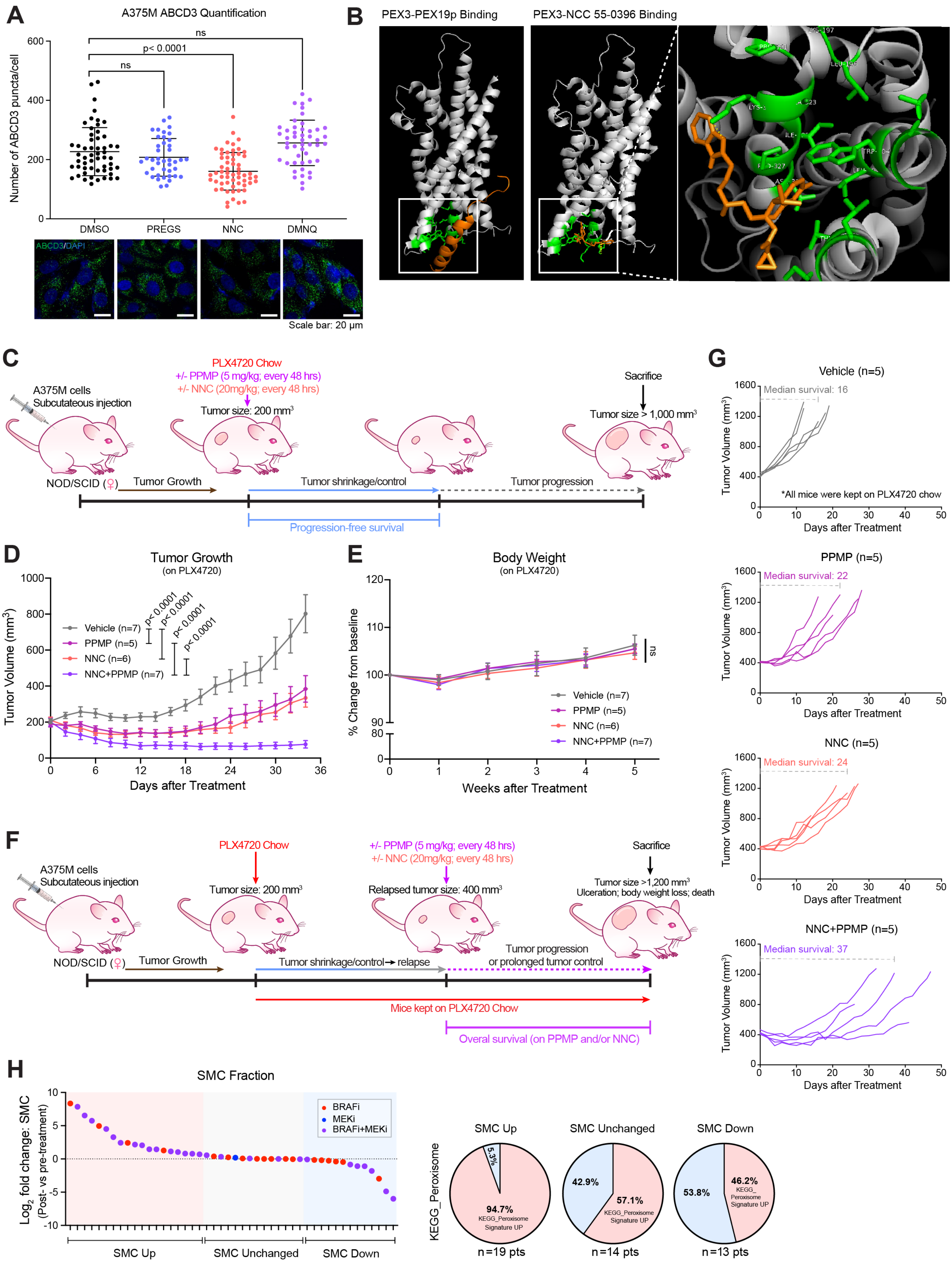
A
A375M ABCD3 Quantification
Number of ABCD3 puncta/cell
ns
p< 0.0001
ns
DMSO
PREGS
NNC
DMNQ
ABCD3/DAPI
Scale bar: 20 μm
B
PEX3-PEX19p Binding
PEX3-NCC 55-0396 Binding
C
PLX4720 Chow
+/- PPMP (5 mg/kg; every 48 hrs)
+/- NNC (20mg/kg; every 48 hrs)
A375M cells
Subcutaneous injection
Tumor size: 200 mm³
Sacrifice
Tumor size > 1,000 mm³
NOD/SCID (♀)
Tumor Growth
Tumor shrinkage/control
Tumor progression
Progression-free survival
D
Tumor Growth
(on PLX4720)
Tumor Volume (mm³)
Vehicle (n=7)
PPMP (n=5)
NNC (n=6)
NNC+PPMP (n=7)
p< 0.0001
p< 0.0001
p< 0.0001
p< 0.0001
Days after Treatment
E
Body Weight
(on PLX4720)
% Change from baseline
ns
Vehicle (n=7)
PPMP (n=5)
NNC (n=6)
NNC+PPMP (n=7)
Weeks after Treatment
F
PLX4720 Chow
+/- PPMP (5 mg/kg; every 48 hrs)
+/- NNC (20mg/kg; every 48 hrs)
Sacrifice
A375M cells
Subcutaneous injection
Tumor size: 200 mm³
Relapsed tumor size: 400 mm³
Tumor size >1,200 mm³
Ulceration; body weight loss; death
NOD/SCID (♀)
Tumor Growth
Tumor shrinkage/control→ relapse
Tumor progression
or prolonged tumor control
Mice kept on PLX4720 Chow
Overal survival (on PPMP and/or NNC)
G
Vehicle (n=5)
Median survival: 16
*All mice were kept on PLX4720 chow
PPMP (n=5)
Median survival: 22
NNC (n=5)
Median survival: 24
NNC+PPMP (n=5)
Median survival: 37
Tumor Volume (mm³)
Days after Treatment
H
SMC Fraction
Log2 fold change: SMC
(Post- vs pre-treatment)
BRAFi
MEKi
BRAFi+MEKi
SMC Up
SMC Unchanged
SMC Down
KEGG_Peroxisome
5.3%
94.7%
KEGG_Peroxisome Signature UP
42.9%
57.1%
KEGG_Peroxisome Signature UP
53.8%
46.2%
KEGG_Peroxisome Signature UP
n=19 pts
n=14 pts
n=13 pts

**Figure S8. PEX3-PEX19 binding inhibitor NNC 55-0396 demonstrates anti-tumoral activity in preclinical melanoma models.**

**(A)** Number of ABCD3 puncta in A375M cells treated with DMSO control, pregnenolone sulfate (PREGS, 10μM), NNC 55-0396 (NNC, 4μM), or 2,3-Dimethoxy-1,4-naphthoquinone (DMNQ, 5μM) for 72 hours. Representative IF staining for ABCD3 (green) and DAPI nuclear stain (blue) are presented (data pooled from n=3 independent experiments). One-way ANOVA. **(B)** Structural analysis showing human PEX19-binding site on PEX3 (left) and predicted binding site of NNC on human PEX3 protein in close proximity to the PEX19-binding site (right). Figures were generated using AutoDock. **(C, F)** Schematic of the experimental design, related to Figure 7F-7I. **(D)** Tumor growth curve (related to Figure 7F-7H) and **(E)** relative weight change (% initial body weight prior to treatment) of mice bearing A375M-derived melanomas receiving indicated treatments. **(G)** Individual growth of A375M-derived melanomas in mouse receiving indicated treatments, related to Figure 7I. Mice were treated with vehicle, NNC, PPMP, or NNC+PPMP after relapsed (PLX4720-resistant) tumor reached a volume of 400 $mm^3$. Median survival is indicated in each graph. All mice were kept on PLX4720 chow after PLX4720 treatment initiated when individual tumor first reached a volume of 200 $mm^3$ (n=5 per group). **(H)** Left: $Log_2$ fold change of SMC abundance in samples from each patient collected post- versus pre-treatment with indicated MAPK inhibitors. A 1.5-fold increase is considered as "SMC up", and a 5% decrease is considered as "SMC down". Right: Pie charts showing percentage of patients in each group with increased or decreased transcript levels of peroxisome-related genes (KEGG_Peroxisome), related to Figure 1A. Total n=46 patients. **(D, E, H, I)** Two-way ANOVA. **(A, I)** Data represented as mean ± SD. **(D, E)** Data represented as mean ± SEM.

**Supplementary Table 1.** Detailed information of human cell lines, culture conditions, treatment, and experimental timeline.

| Cell name | Cultured in | Treatment | Timeline |
|---|---|---|---|
| A375M | DMEM | Vemu (2.5 μM) | Day 1: seed + siRNA transfection<br>Day 3: treatment<br>Day 4: harvest (24h treatment) |
| | | Vemu (2.5 μM)<br>PPMP (7.5 μM) | Day 1: seed + siRNA transfection<br>Day 3: treatment<br>Day 4: harvest (24h treatment) |
| 1205Lu | RPMI | Vemu (5 μM)<br>Cobi (50 nM) | Day 1: seed + siRNA transfection<br>Day 3: treatment<br>Day 4: harvest (24h treatment) |
| | | Vemu (5 μM)<br>Cobi (50 nM)<br>PPMP (10 μM) | Day 1: seed + siRNA transfection<br>Day 3: treatment<br>Day 4: harvest (24h treatment) |
| WM3406 | RPMI<br>1x GlutaMax | Cobi (100 nM) | Day 1: seed + siRNA transfection<br>Day 2: treatment<br>Day 4: harvest (48h treatment) |
| | | Cobi (100 nM)<br>PPMP (5 μM) | Day 1: seed + siRNA transfection<br>Day 2: treatment<br>Day 4: harvest (48h treatment) |
| MeWo | DMEM | Cobi (100 nM) | Day 1: seed + siRNA transfection<br>Day 2: treatment<br>Day 4: harvest (48h treatment) |
| | | Cobi (100 nM)<br>PPMP (5 μM) | Day 1: seed + siRNA transfection<br>Day 2: treatment<br>Day 4: harvest (48h treatment) |
| MelST | DMEM | Vemu (5 μM)<br>Cobi (100 nM) | Day 1: seed + siRNA transfection<br>Day 2: treatment<br>Day 4: harvest (48h treatment) |
| A375 VSR | DMEM<br>Vemu (2.5 μM) | Vemu (2.5 μM)<br>PPMP (7.5 μM) | Day 1: seed + siRNA transfection<br>Day 2: treatment<br>Day 4: harvest (48h treatment) |
| 1205Lu VCDR | RPMI<br>Vemu (5 μM)<br>Cobi (50 nM) | Vemu (5 μM)<br>Cobi (50 nM)<br>PPMP (5 μM) | Day 1: seed + siRNA transfection<br>Day 2: treatment<br>Day 4: harvest (48h treatment) |
| WM164 VSR | DMEM<br>Vemu (1 μM) | Vemu (1 μM)<br>PPMP (10 μM) | Day 1: seed + siRNA transfection<br>Day 2: treatment<br>Day 4: harvest (48h treatment) |
| SK-Mel-28 VSR | DMEM<br>Vemu (2.5 μM) | Vemu (2.5 μM)<br>PPMP (10 μM) | Day 1: seed + siRNA transfection<br>Day 2: treatment<br>Day 4: harvest (48h treatment) |
| SK-Mel-28<br>VCDR | DMEM<br>Vemu (2.5 μM)<br>Cobi (100 nM) | Vemu (2.5 μM)<br>Cobi (100 nM)<br>PPMP (5 μM) | Day 1: seed + siRNA transfection<br>Day 2: treatment<br>Day 4: harvest (48h treatment) |
| MeWo CSR | DMEM<br>Cobi (400 nM) | Cobi (400 nM)<br>PPMP (5μM) | Day 1: seed + siRNA transfection<br>Day 2: add Cobi<br>Day 3: add PPMP<br>Day 5: harvest (72h Cobi+48h PPMP) |

* All cells were cultured in the indicated medium supplemented with 10% FBS and 1x Pen/Strep.

**Supplementary Table 2.** siRNAs.

| Gene name | siRNA | Duplex Sequence (5’-3’) |
|---|---|---|
| ***PEX3 (human)*** | *siPEX3* | rCrGrGrArCrArGrArUrCrCrArUrUrCrArGrUrUrUrGrCrAGT |
| | | rArCrUrGrCrArArArCrUrGrArArUrGrGrArUrCrUrGrUrCrCrGrUrUr |
| | *siPEX3-2* | rGrArUrCrUrGrArArGrArUrAArUrArArGrUrUrUrCrArCAA |
| | | rUrUrGrUrGrArArArCrUrUrArUrUrArUrCrUrUrCrArGrArUrCrCrU |
| ***PEX19 (human)*** | *siPEX19* | rGrUrGrArArCrArGrUrGrUrCrUrGrArUrCrArUrGrUrGrAAA |
| | | rUrUrUrCrArCrArUrGrArUrCrArGrArCrArCrUrGrUrUrCrArCrCrA |
| ***UGCG (human)*** | *siUGCG-1* | rCrUrUrCrArCrArUrCrCrArArGrArUrArCrUrArUrArUrCTC |
| | | rGrArGrArUrArUrArGrUrArUrCrUrUrGrGrArUrGrUrGrArArGrUrU |
| | *siUGCG-2* | rGrCrUrUrUrGrUrGrArCrUrGrUrArUrArUrArArArGrGrAAA |
| | | rUrUrUrCrCrUrUrUrArUrArUrArCrArGrUrCrArCrArArArGrCrUrG |
| ***Ugcg (mouse)*** | *siUgcg-1* | rArGrArArUrGrUrArArUrUrUrCrArUrGrArUrArCrArArGTA |
| | | rUrArCrUrUrGrUrArUrCrArUrGrArArArUrUrArCrArUrUrCrUrUrA |
| | *siUgcg-2* | rGrUrArCrArUrUrGrCrUrGrArArGrArUrUrArCrUrUrUrATG |
| | | rCrArUrArArArGrUrArArUrCrUrUrCrArGrCrArArUrGrUrArCrUrG |
| ***Gba (mouse)*** | *siGba-1* | rGrGrUrUrCrCrArArGrArGrCrUrArUrGrArUrArUrCrUrGTC |
| | | rGrArCrArGrArUrArUrCrArUrArGrCrUrCrUrUrGrGrArArCrCrGrA |
| | *siGba-2* | rGrUrGrArArGrCrUrArCrUrCrArUrGrCrUrArGrArUrGrACC |
| | | rGrGrUrCrArUrCrUrArGrCrArUrGrArGrUrArGrCrUrUrCrArCrArU |
| ***Negative control*** | *siCtrl* | N/A (Proprietary, AllStar Neg. Control siRNA, QIAGEN #1027218) |

**Supplementary Table 3.** Detailed information of the primary antibodies used for western blotting, immunohistochemistry, and immunofluorescence staining.

| Target | Antibody full name | Source & Catalog # | Experiment |
|---|---|---|---|
| PEX3 | Anti-PEX3 antibody produced in rabbit | Sigma-Aldrich, HPA042830 | WB, IF (human) |
| PEX3 | PEX3 Polyclonal Antibody | ThermoFisher, PA5-115740 | WB (murine) |
| PEX19 | PEX19 Polyclonal antibody | Proteintech, 14713-1-AP | WB |
| PEX16 | PEX16 Polyclonal antibody | Proteintech, 14816-1-AP | WB |
| UGCG | UGCG Antibody (1E5) | Novus Biologicals, H00007357-M03 | WB |
| UGCG | UGCG Polyclonal antibody | Proteintech, 12869-1-AP | IF |
| AGPS | Anti-AGPS antibody | Abcam, ab236621 | WB |
| CD36 | Brilliant Violet 421™ anti-human CD36 Antibody | BioLegend, 336229 | Flow, IF |
| ABCD3 | Anti-PMP70 antibody [CL2524] | Abcam, ab211533 | IF |
| ABCD3 | Rabbit polyclonal anti-PMP70 antibody | Abcam, ab3421 | IF |
| Myc-Tag | Myc-Tag (9B11) Mouse mAb | Cell Signaling #2276 | IP |
| GAPDH | GAPDH (14C10) Rabbit mAb | Cell Signaling #2118 | WB |
| α-Actinin | Anti-α-actinin Antibody (H-2) | Santa Cruz, sc-17829 | WB |
| β-Actin | Monoclonal Anti-β-Actin antibody (clone AC-15) | Sigma-Aldrich #A5441 | WB |

**Supplementary Table 4.** RT-qPCR primers.

| Gene name | Primer | Sequence (5'-3') |
|---|---|---|
| ***AGPS*** | Fwd | GTGACCCACTGACCGTATTT |
| | Rev | GCCATTGCTTCCGTAACTTG |
| ***PEX1*** | Fwd | CCTGTGTGCTACAAGTAGTCTG |
| | Rev | GGAATCCAGACTTTCCCAAGA |
| ***SCP2*** **(N terminal)** | Fwd | AACTGTGCTACTGGTTCTACTG |
| | Rev | GGCTTCCCTTACTCATCTTCTC |
| ***UGCG*** | Fwd | GGATCAAGCAGGAGGACTTATAG |
| | Rev | CTTGAGTGGACATTGCAAACC |
| ***Rplp0 (m36B4)*** | Fwd | TCATCCAGCAGGTGTTTGACA |
| | Rev | GGCACCGAGGCAACAGTT |
| ***Actb*** | Fwd | GGCTGTATTCCCCTCCATCG |
| | Rev | CCAGTTGGTAACAATGCCATGT |
